# An EEG Signature of MCH Neuron Activities Predicts Cocaine Seeking

**DOI:** 10.1101/2024.03.27.586887

**Authors:** Yao Wang, Danyang Li, Joseph Widjaja, Rong Guo, Li Cai, Rongzhen Yan, Sahin Ozsoy, Giancarlo Allocca, Jidong Fang, Yan Dong, George C. Tseng, Chengcheng Huang, Yanhua H. Huang

**Affiliations:** Departments of Psychiatry, University of Pittsburgh, Pittsburgh, PA 15219; Neuroscience, University of Pittsburgh, Pittsburgh, PA 15260; Biostatistics, University of Pittsburgh, Pittsburgh, PA 15213; North Allegheny High School, Pittsburgh, PA 15237; Somnivore Pty. Ltd., Bacchus Marsh, VIC, Australia 3340; Department of Pharmacology and Therapeutics, The University of Melbourne, VIC, Australia 3010; The Florey Institute of Neuroscience and Mental Health, The University of Melbourne, VIC, Australia; Department of Psychiatry and Behavioral Health, Penn State College of Medicine, Hershey, PA 17033

**Keywords:** EEG, MCH, REM sleep, cocaine, biomarker

## Abstract

**Background:** Identifying biomarkers that predict substance use disorder (SUD) propensity may better strategize anti-addiction treatment. The melanin-concentrating hormone (MCH) neurons in the lateral hypothalamus (LH) critically mediates interactions between sleep and substance use; however, their activities are largely obscured in surface electroencephalogram (EEG) measures, hindering the development of biomarkers.

**Methods:** Surface EEG signals and real-time Ca^2+^ activities of LH MCH neurons (Ca^2+^_MCH_) were simultaneously recorded in male and female adult rats. Mathematical modeling and machine learning were then applied to predict Ca^2+^_MCH_ using EEG derivatives. The robustness of the predictions was tested across sex and treatment conditions. Finally, features extracted from the EEG-predicted Ca^2+^_MCH_ either before or after cocaine experience were used to predict future drug-seeking behaviors.

**Results:** An EEG waveform derivative – a modified theta-to-delta ratio (EEG Ratio) – accurately tracks real-time Ca^2+^_MCH_ in rats. The prediction was robust during rapid eye movement sleep (REMS), persisted through REMS manipulations, wakefulness, circadian phases, and was consistent across sex. Moreover, cocaine self-administration and long-term withdrawal altered EEG Ratio suggesting shortening and circadian redistribution of synchronous MCH neuron activities. In addition, features of EEG Ratio indicative of prolonged synchronous MCH neuron activities predicted lower subsequent cocaine seeking. EEG Ratio also exhibited advantages over conventional REMS measures for the predictions.

**Conclusions:** The identified EEG Ratio may serve as a non-invasive measure for assessing MCH neuron activities *in vivo* and evaluating REMS; it may also serve as a potential biomarker predicting drug use propensity.

## Introduction

Identifying biomarkers that predict future drug use propensity may open new avenues for individualized medicine in preventing and treating substance use disorders (SUDs) (1, 2). Lateral hypothalamus (LH), a phylogenetically conserved central processor, controls vital body functions and regulates reward and motivation through a variety of molecularly distinct cell types (3–6). The melanin-concentrating hormone (MCH) neurons are deeply embedded in the LH and zona incerta, projecting throughout the brain to regulate sleep, energy balance, emotion, reward, and motivation (7–16). MCH neuron activity is an integral component of rapid eye movement sleep (REMS) (10, 17), and REMS powerfully regulates the formation and modification of emotional memories (18–20). Following cocaine self-administration and withdrawal, MCH neurons in the rat exhibit reduced membrane excitability and decreased glutamate receptor activities (21), whereas enhancing MCH neuron activities in sleep decreases relapse-like behaviors, accompanied by reversal of cocaine-induced cellular changes in the reward circuit (22). Thus, MCH neuron activities may provide important functional measures for sleep, emotion regulation, and drug relapse propensity. However, MCH neuron activities can only be monitored through invasive and low-throughput *in vivo* recordings combined with genetic labeling of these neurons (17, 18, 22, 23).

Surface electroencephalogram (EEG), or scalp EEG, depicts the temporal dynamics of whole-brain electric activities, though predominantly cortical activities, due to close proximity (24). Whether and to what extent subcortical activities are reflected in surface EEG are still being explored. Through simulations and high-density EEG recordings, it was shown possible to derive subcortical activities such as in the thalamus (25, 26). However, it is not known whether activities in deep structures such as the lateral hypothalamus may be deduced from surface EEG signals.

Here, we applied mathematical modeling and machine learning to surface EEG signals to predict real-time Ca^2+^ activities of LH MCH neurons (Ca^2+^_MCH_) *in vivo*. We tested the robustness of the predictions across vigilance states, circadian phases, REMS manipulations, cocaine self-administration and withdrawal, and sex. We evaluated whether the EEG Ratio may serve as a non-invasive measure for assessing MCH neuron activities *in vivo* and, therefore, functional aspects of REMS. Furthermore, we used features extracted from the EEG Ratio either before or after cocaine experience to predict future drug-seeking behaviors.

## Methods and Materials

### Animals

Male and female Sprague Dawley rats were ∼postnatal day (PD) 70 at the start of recordings. Both male and female rats were used to characterize EEG–Ca^2+^_MCH_ correlations (Figs. 1-4), and male rats were further tested with cocaine self-administration protocols (Figs. 5-7). Rats were singly housed under a 12-h reverse light/dark cycle with controlled temperature (21-22°C) and humidity (60±5%). All experiments were performed in accordance with University of Pittsburgh IACUC-approved protocols. Cocaine self-administration training and incubation test (22, 27–29).

**Figure 1.**
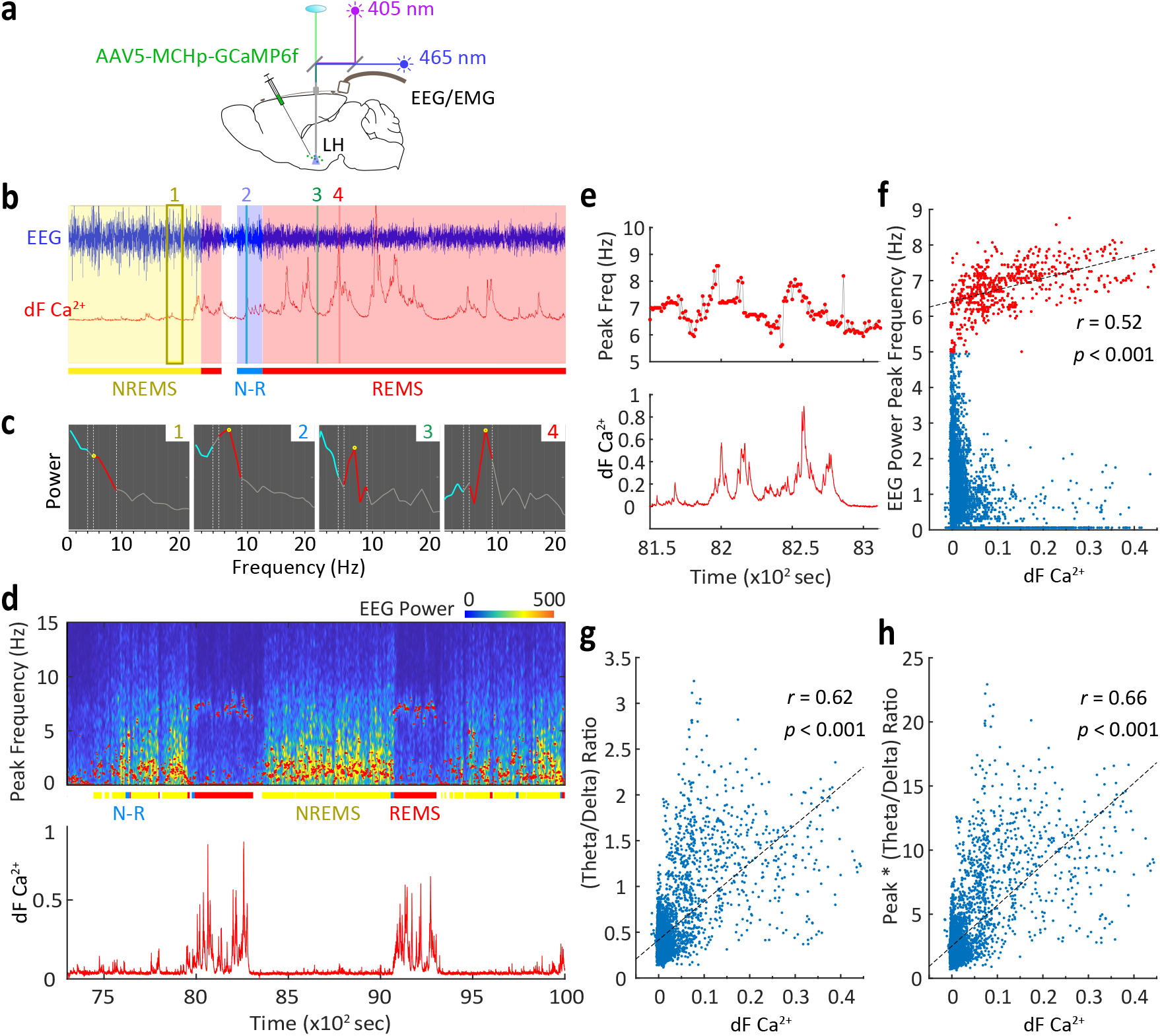
Ca^2+^_MCH_ transients correlate with coordinated changes in EEG *θ* and *δ* activities in sleep. **a** EEG and fiber photometry dual recording diagram. **b** Ca^2+^_MCH_ transients occurred predominantly during REMS. **c** Ca^2+^_MCH_ peaks were accompanied by an increase in *θ* (*red*) and a decrease in *δ* (*blue*) magnitude, with a shift in the peak of *θ* frequency (*yellow circle*) towards higher *θ* frequency. **c** EEG spectrogram aligned with Ca^2+^_MCH_ over NREM and REM episodes, showing increase in t and decrease in d activities during Ca^2+^_MCH_ responses. *Red* dots in *upper* panel represent peak frequencies of EEG power spectrum (0-20 Hz) at 1-sec resolution. **e** Peak frequencies of *θ* activities during a REMS episode from **d**. **f** Peak frequencies of *θ* activities showed a positive correlation with Ca^2+^_MCH_ amplitude over a 2-h recording period. **g** *θ/δ* ratio showed a positive correlation with Ca^2+^_MCH_ amplitude over the same 2-h recording period. **h** *θ* peak x (*θ/δ* ratio) showed a positive correlation with Ca^2+^_MCH_ amplitude over the same 2-h recording period, with an improved *r*-value.

**Figure 2.**
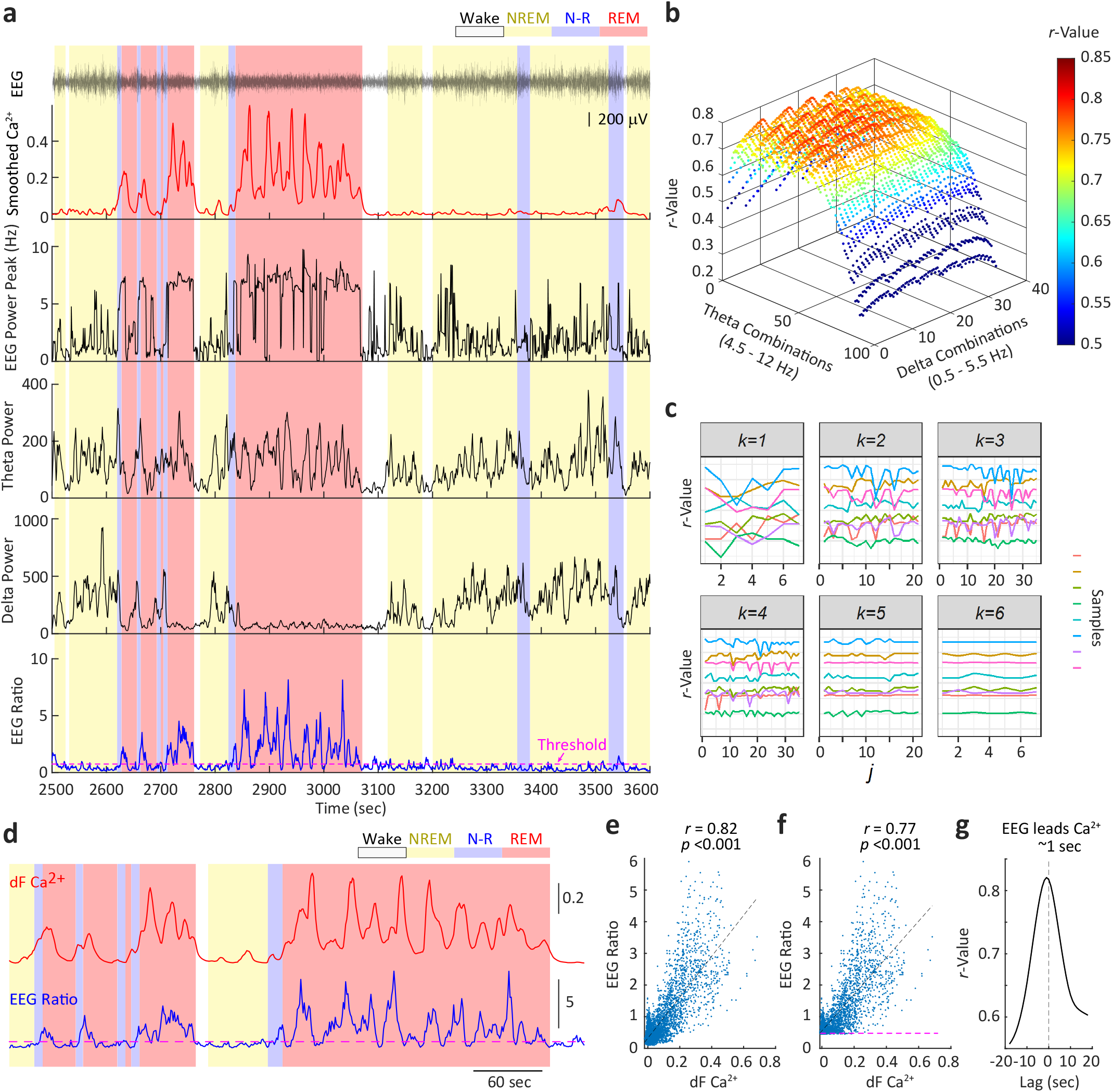
Ca^2+^_MCH_ is correlated with EEG Ratio: empirical formula and cross-validation. **a** Example Ca^2+^_MCH_ aligned to EEG and individual EEG features calculated over 10-sec sliding windows at 1-sec steps. Ca^2+^_MCH_ was smoothed using the same sliding window. EEG Ratio = (Power *θ*/Power *δ*) x (F_peak_/7_Hz_); threshold (*pink dotted line*) = amplitude at 60% dwelling time over 2-h recordings. **b** *r*-values for EEG Ratio – Ca^2+^_MCH_ correlations from an example 2-h recording, which were calculated using 36 x 91 combinations of extended *θ* and *δ* ranges at 0.5 Hz increments for optimization. **c** Cross-validation for training sample size *k* = 1,…, 7. When *k ≥* 6, the performance became stable. **d** Example EEG Ratio calculated using the optimized formula: EEG Ratio = (Power *θ*_6.5-9.5 Hz_ / Power *δ*_1-5.5 Hz_) x (F_peak_ / 7 Hz), and aligned to Ca^2+^_MCH_. **e** EEG ratio – Ca^2+^_MCH_ amplitude correlation over a 2-h recording in an example rat. **f** Supra-threshold EEG Ratio – Ca^2+^_MCH_ amplitude correlation over the same 2-h recordings in the example rat. **g** *r*-values at different EEG Ratio – Ca^2+^_MCH_ time lags in the example rat.

**Figure 3.**
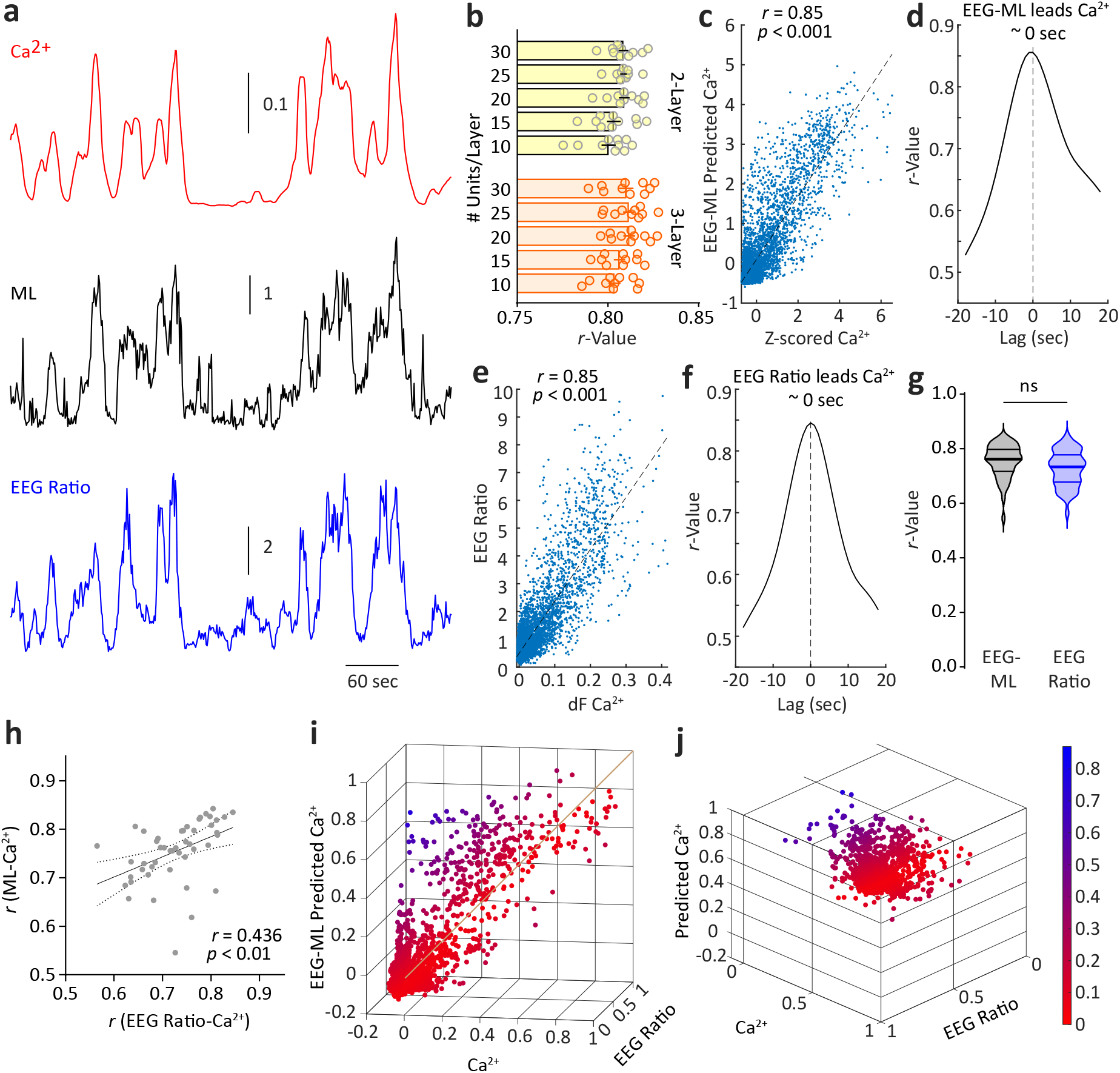
Ca^2+^_MCH_ can be predicted from EEG signals using machine-learning (ML). **a** Example Ca^2+^_MCH_ over time aligned with ML-predicted, amplitude-normalized (i.e. Z-scored), Ca^2+^_MCH_, and the corresponding empirically calculated EEG Ratio. **b** ML using *r*-values to optimize # of layers and # of units per layer for the multi-layer perceptron network, with 10-fold cross-validation (shown in *circles*) for each condition. These networks yielded similar performances on the pilot data (layer x unit interaction: F_4, 90_=0.020, *p*=0.999; two-way ANOVA), therefore we chose the one with the best average performance: networks with 3 hidden layers and 20 units/layer. **c** EEG-ML– Ca^2+^_MCH_ amplitude correlation over a 2-h recording in the example rat using an optimized multi-layer perceptron network with 3 hidden layers and 20 units/layer. **d** *r*-values at different EEG-ML – Ca^2+^_MCH_ time lags in the example rat. **e** EEG ratio – Ca^2+^_MCH_ amplitude correlation over a 2-h recording in the example rat. **f** *r*-values at different EEG Ratio – Ca^2+^_MCH_ time lags in the example rat. **g** Among the testing set (n=36 new 2-h recordings), EEG-ML versus EEG Ratio performance was not significantly different (*t*_35_=0.838, *p*=0.408, n=36 rats each, male and female, paired *t*-test). ns = not significant. **h** The *r*-values for ML-Ca^2+^_MCH_ predictions versus EEG Ratio Ca^2+^_MCH_ predictions were correlated. Each *circle* represents one rat and/or condition. n=47 rats and/or conditions, male and female. **i** Normalized Ca^2+^_MCH_ (*x*-axis), EEG Ratio (*y*-axis), and ML-predicted Ca^2+^ (*z*-axis) triplet-pairs were plotted within a 1×1×1 cube, using data from the example rat. Each dot represents a 10-sec sample from the 2-h recording. Color gradient represents the distance to the diagonal line (*x*=*y*=*z*). **j** Same plot as **i** viewed along the diagonal line.

**Figure 4.**
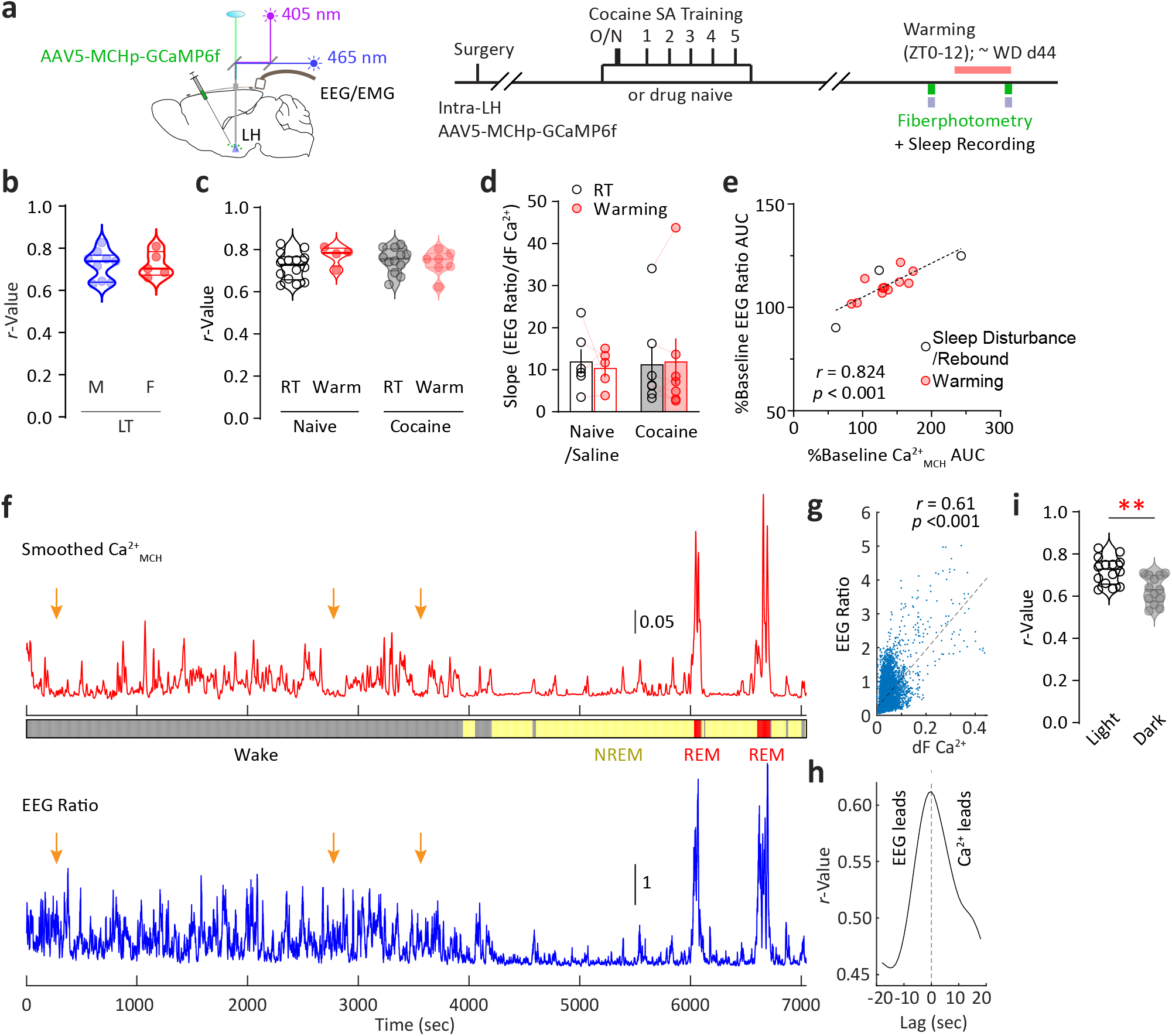
EEG Ratio – Ca^2+^_MCH_ correlations across sex, treatment conditions, and light/dark phases. **a** EEG and fiber photometry dual recording diagram and timeline. **b** EEG Ratio – Ca^2+^_MCH_ correlation *r*-values were similar in males (n=9) and females (n=5) (*t*_12_=0.112, *p*=0.913, *t*-test). **c** EEG Ratio – Ca^2+^_MCH_ correlations persisted under REMS manipulations by environmental warming in naïve/saline-treated rats or following cocaine experience (interaction: F_1, 33_=1.913, *p*=0.176, two-way ANOVA). **d** The slope of the EEG Ratio – Ca^2+^_MCH_ plot was not altered following cocaine exposure or REMS manipulations by environmental warming (interaction: F_1, 10_=0.612, *p*=0.452, two-way RM ANOVA). **e** %Changes in EEG Ratio versus % changes in Ca^2+^_MCH_ responses under sleep disturbance/rebound, and environmental warming to increase REMS; within-subject comparisons. **f** In the dark phase during explorative behaviors in wakefulness, there was asynchronous Ca^2+^_MCH_ activity accompanied by high-frequency, low-amplitude EEG events. **g**-**h** EEG Ratio-Ca^2+^_MCH_ correlation from the example 2-h recordings. Each *dot* in **g** represents a 10-sec sample from the recording. **i** Overall reduced correlation *r*-values during the randomly sampled dark phase hours compared to light phase (*t*_25_=3.650, *p*<0.01, *t*-test). *arrows* in **f** indicate occasions of EEG events when Ca^2+^_MCH_ was absent. Each *circle* represents one rat. n=5-14 male and female rats.

**Figure 5.**
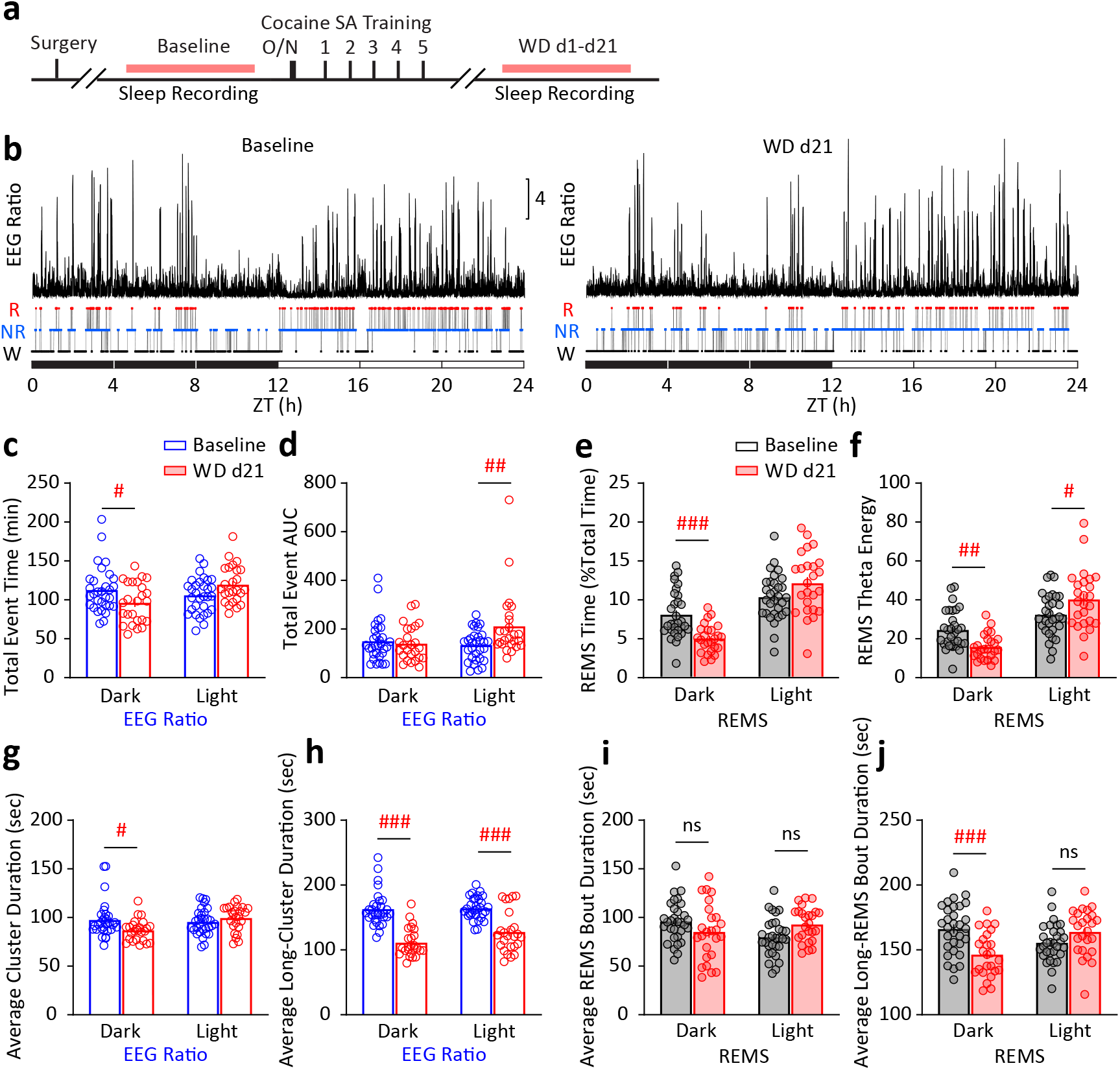
Cocaine-induced long-term changes in EEG Ratio. **a** Cocaine self-administration training and EEG recording timeline. **b** Example 24-h EEG Ratio extracted from baseline sleep before cocaine exposure (*left*) or after 21 days of withdrawal (*right*) from repeated cocaine self-administration (1 overnight + 2-h daily x 5 d, 0.75 mg/kg/infusion, FR1). **c**,**d** The 24-h EEG Ratio exhibited reduced activities in the dark phase and increased activities in the light phase at withdrawal d21, measured by the total event time (**c**) or total AUC (**d**). **c** interaction: F_1, 54_=8.804, *p*<0.01; **d** interaction: F_1, 54_=9.815, *p*<0.01; two-way RM ANOVA with Sidak *post-hoc* test. **e**,**f** Similar changes were observed in REMS, measured by the total REMS time (**e**) or *θ* energy (**f**) during REMS (*θ* energy = *θ* power x duration). **e** interaction: F_1, 54_=12.97, *p*<0.001; **f** interaction: F_1, 54_=13.79, *p*<0.001; two-way RM ANOVA with Sidak *post-hoc* test. **g** Cluster analysis of EEG Ratio showed a decrease in the average cluster durations in the dark phase and an increase in the light phase. **h** Long (>50 sec) EEG event clusters showed a decrease in durations in both dark and light phases. **g** interaction: F_1, 54_=5.053, *p*<0.05; **h** interaction: F_1, 54_=3.458, *p*=0.068; two-way RM ANOVA with Sidak *post-hoc* test. **i** In contrast to **g**, average REMS bout duration did not change in either dark or light phase. **j** In contrast to **h**, average long-bout REMS did not change in light phase. **i** interaction: F_1, 54_=11.67, *p*<0.01; **j** interaction: F_1, 54_=14.99, *p*<0.001; two-way RM ANOVA with Sidak *post-hoc* test. # *p*<0.05, ## *p*<0.01, ### *p*<0.001 *post-hoc* test. Each circle represents a rat. n=25-31 male rats.

### Viral vector

AAV5-MCHp-GCaMP6f was constructed by connecting MCH promoter (MCHp) with GCaMP6f (Addgene #52925), packaged at University of North Carolina Vector Core, and diluted to ∼ 1-5×10^12^ vg/ml working concentration. The expression specificity was validated previously (22).

### Surgeries

Cocaine self-administration surgery, EEG/EMG surgery, AAV stereotaxic injections, and fiberoptic implantations were similar as previously described (21, 22, 27, 30–34), and combined per experimental designs. AAV5-MCHp-GCaMP6f was injected at 1.0 µl/site (0.1 µl/min) into the LH and zona incerta (in mm): AP -2.56, ML +/-0.90, DV -8.10. For *in vivo* fiber photometry, optic fibers (400-µm core) with flat tips were unilaterally implanted directly above zona incerta (AP -2.56, ML +/-1.50, DV -7.50). For EEG/EMG surgery, two custom-made wire EMG electrodes were knotted into the nuchal neck muscle, and two pairs of screw-mounted wire EEG electrodes (E363/20 P1 technology, Roanoke, VA) were installed contralaterally through the skull over the parietal/occipital and frontal cortices (AP +4, ML +/-2.5; AP -6, ML +/-3). All rats were single-housed post-surgery and during subsequent experimentation. Rats started recordings or behavioral training ∼2 weeks after surgeries.

### Combined sleep and fiber photometry recordings and initial data processing

Dual EEG and fiber photometry recordings were achieved through concentric configuration of the two recording cables, similar to previously described (22). Rats habituated in the recording chamber for 3 days after being tethered before recordings. EEG (0.1–30 Hz) and EMG (30 Hz–3 kHz) signals were amplified using Grass model 15LT bipolar amplifiers (Grass Technologies, West Warwick, RI) and converted by an analog-digital converter (Kissei Comtec). All signals were digitized at 128 Hz and collected using Vital Recorder software (Kissei Comtec, Nagano, Japan) or SleepMaster (Biosoft Studio, PA, USA). Sleep recordings were analyzed using SleepMaster (Biosoft Studio, PA, USA), Somnivore 2.0 (Somnivore Pty Ltd, Australia) (35), and MATLAB R2019a custom-written scripts.

Dual-channel fiber photometry was performed to monitor GCaMP6f signals (465 nm light) with isosbestic control signals independent of Ca^2+^ activity (405 nm light; at ∼30 µW each; LEDs, Doric Lenses) as described previously (22). GCaMP6f fluorescence (F_465_) and isosbestic signals (F_405_) were collected by a femtowatt photoreceiver (New Focus 2151, Newport), amplified, and A-D converted using the RZ5P processor (Tucker David Technologies, USA). GCaMP6f recordings lasted for 2 h each time and were TTL time-stamped by the sleep recording program. Raw photometry data (F_405_, F_465_) were recorded using Synapse software (Tucker David Technologies, USA) and analyzed using MATLAB R2019a custom-written scripts. F_405_ and F_465_ were aligned to 1-sec epoch sleep scores (Somnivore 2.0) using TTL time stamps. Detrending and normalization of each light-on period was calculated by dF = F_465_ / Median_465_ – F_405_ / Median_405_.

### Cocaine self-administration training and testing

Cocaine self-administration training (0.75 mg/kg/infusion, 1 overnight + 5 d x 2 h/d) was similar as described previously (21, 22, 27). It was conducted during the dark phase in operant-conditioning chambers (Med Associates, VT). Rats were trained using a fixed ratio (FR) 1 reinforcement schedule (0.75 mg/kg over 6 s per infusion, 20 s time-out). Nose-pokes in the inactive hole had no reinforcement consequences. Rats that received at least 20 cocaine infusions during the overnight session were subject to a subsequent 5-day training (2 h daily). On withdrawal d1 and d45, rats were tested for 30 min for cue-induced cocaine seeking in the same boxes without receiving cocaine infusions. Cocaine HCl (provided by NIDA Drug Supply Program) was dissolved in 0.9% NaCl saline.

### Warming

The sleep recording cage was placed on top of a rectangular electrical heating pad (TK-SHM, Seedfactor) with feed-back control set to 31.5-34.5°C during ZT1-11 (10 h in the light period). Details see (22).

### EEG Ratio calculation, optimization, and cross-validation

EEG Ratio calculation and optimization were performed using MATLAB R2019a custom-written scripts. EEG signals underwent Fast Fourier Transformation (FFT) using 0.5 Hz frequency bins (0.5-50 Hz). EEG Ratio base formula was empirically defined as the theta (*θ*, 0.5-4 Hz)-to-delta (*δ*, 5-9 Hz) power ratio multiplied by peak frequency (F_peak_) relative to the mid-*θ* frequency (7 Hz), i.e. EEG Ratio base formula = (Power *θ*/Power *δ*) x (F_peak_/7_Hz_). *θ* and *δ* ranges were then tested systematically at 0.5 Hz increments at both lower and higher boundaries (*θ:* 4.5-12 Hz*; δ:* 0.5-5.5 Hz). The smoothing window was tested at 1 – 10 sec at 1 sec increment. The linear regression *r*-values for EEG Ratio – Ca^2+^_MCH_ amplitude correlation were then calculated for a 2-h recording in each rat using all 91 x 36 combinations of the *θ* and *δ* ranges and 10 different smoothing windows. The *r*-value distributions were then analyzed for optimal ranges for *θ*, *δ*, and the smoothing window for each individual rat. The optimized EEG Ratio = (Power *θ*_6.5-9.5 Hz_/Power *δ*_1-5.5 Hz_) x (F_peak_/7_Hz_), using 10-sec sliding window at 1 sec steps.

Next, cross-validation was performed using R software version 4.2.1 to evaluate the ability of using the optimized EEG ratio to predict Ca^2+^_MCH_ in new testing samples, quantified by the correlation *r*-value of EEG ratio and Ca^2+^_MCH_. In each iteration, *k* out of 8 rats were subsampled as training set and the remaining rats as test set. The optimized *θ* and *δ* power ranges were selected that generated the highest averaged correlation *r*-values. The optimized *θ* and *δ* power ranges were then used to validate correlation in the test set. The analysis was repeated for *k* = 1,…, 7. For a given *k*, each rat treated in the test set has 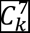 unique training set from the remaining 7 rats and the cross-validated *r*-values were compared.

### Machine learning-based prediction of Ca2+MCH using EEG signals

Machine-learning was performed using MATLAB R2019a custom-written scripts. We applied a multi-layer perceptron (MLP) network with a rectified linear activation function and trained with stochastic gradient descent with momentum method (36, 37). The time series of EEG power and Ca^2+^_MCH_ signals were Z-scored for each animal, and broken into partially overlapping time windows (10 sec each), where each time window was treated as a data sample. The input to the MLP network is the power of the EEG signal across frequency channels (0 – 20 Hz) during a time window, and the output of the network was optimized to match the mean magnitude of Ca^2+^_MCH_ during the same time window. The MLP network was optimized to be fully connected with 3 hidden layers of 20 units each. We used data from 6 pilot rats (2 h each) for training the MLP network with 10-fold cross-validation, then evaluated generalizability in ∼ 36 additional independent 2-h recordings from 23 rats.

### Statistical analysis for grouped data

Rats were randomly assigned to all experimental groups. Group sizes were determined based on power analyses using preliminary estimates of variance with the goal of achieving 80% power to observe differences at α=0.05. No data points were excluded unless specified in the experimental procedure. All data were analyzed blind to treatment groups. Statistical analysis was performed using Prism GraphPad9. Normal distribution was assumed for all statistics based on visual inspection and previous experience, and was not formally tested (38). Homoscedasticity (equality of variance) was tested using Fisher’s *F* test (between 2 groups) or Levene’s test (3 or more groups) prior to *t* test or ANOVA test respectively (*F* and Levene’s testing results not shown). All tests were two-sided. *p* < 0.05 was considered statistically significant. All rats were coded for sleep scoring, and then decoded for data compiling. All results are presented as mean± SEM unless specified otherwise.

### Assessing prediction

The ability of using EEG features or REMS features to predict subsequent cocaine intake was assessed using R software version 4.2.1. by applying leave-one-out cross-validation. Using baseline data, simple linear regression was performed for each feature leaving one sample out of the training data as the validation data. When outcome was treated as a continuous variable, cross-validated *r* was used to evaluate the predictive performance. When outcome was treated ordinally, it was polychotomized into 3 levels: low (<25%), median (25-75%), and high (>75%), and the predictive performance was evaluated by Somers’ D between the predicted outcomes and true outcomes, and a cross-validated root-mean-square error (rMSE) standardized by the raw rMSE without prediction.

## Results

### Ca^2+^_MCH_ transients correlate with coordinated changes in EEG θ and δ activities in sleep

Simultaneous scalp EEG and fiber photometry recordings of MCH neurons (Fig. 1a) were performed during Zeitgeber time (ZT) 6-8, when REMS occurs frequently (22). Ca^2+^_MCH_ signal F_465_ was normalized to the isosbestic control signal F_405_ and aligned with EEG spectrogram at 1-sec epochs (Methods; Fig. 1b). Ca^2+^_MCH_ peaks were preferentially accompanied by an increase in *θ* (5-9 Hz) and a decrease in *δ* (0.5-4 Hz) power, with a shift in the peak frequency towards higher *θ* frequency (Fig. 1c,d). Moreover, *θ* peak frequency was positively correlated with Ca^2+^_MCH_ levels (Fig. 1e,f). Additionally, although isolated *θ* and *δ* powers were not correlated with Ca^2+^_MCH_ (data not shown), the *θ*/*δ* ratio was positively correlated with Ca^2+^_MCH_ (Fig. 1g). This was consistent with the notion that the overall *θ*/*δ* ratio is elevated during REMS (39, 40), and that MCH neuron activities also predominantly occur during REMS (10, 17). Furthermore, combining the two components by multiplication (*θ* peak frequency x *θ*/*δ* ratio) yielded higher correlations with Ca^2+^_MCH_ (Fig. 1h). These results prompted us to further optimize the Ca^2+^_MCH_-EEG correlations and to test for generalizability.

### EEG Ratio calculated based on empirical formula correlates with Ca^2+^_MCH_

The EEG spectrum was calculated using a 10-sec sliding window at 1 sec increments, and the EEG Ratio was empirically defined as the *θ*-to-*δ* power ratio multiplied by the shift of peak frequency (F_peak_) relative to the mid-*θ* frequency (∼7 Hz), i.e., EEG Ratio = (Power *θ*/Power *δ*) x (F_peak_/7_Hz_) (Fig. 2a). The mid-*θ* frequency 7 Hz was chosen arbitrarily to reference the *θ* peak shift and did not impact the correlation *r*-values (defined below). For each set of 2-h recordings, the sec-by-sec EEG Ratio – Ca^2+^_MCH_ amplitude pairs were fit to a linear regression, and the correlation *r*-value was calculated and used as an index for further improving EEG Ratio calculations. The *θ* and *δ* frequency bands, as well as the smoothing window durations were thus optimized and cross-validated (Methods) among the pilot 8 male rats (Fig. 2b,c, Fig. S1).

As shown in Fig. 2b, multiple combinations of *θ* and *δ* frequency bands could reach the highest *r*-values. We then compared using a single optimal parameter set versus multiple top-performing parameter sets, and found that using multiple sets only mildly reduced the variability while considerably reduced the prediction ability, particularly when the training sample size became larger (Fig. S1). Thus, we used single optimal parameter set for simplicity and better averaged performance, i.e. optimized *θ* = 6.5-9.5 Hz, *δ* = 1-5.5 Hz.

As such, we defined the final optimized EEG Ratio = (Power *θ*_6.5-9.5 Hz_ / Power *δ*_1-5.5 Hz_) x (F_peak_ / 7 Hz), calculated over 10-sec smooth-windows and 1-sec slide-steps. Using this formula, the EEG Ratio showed high positive correlations with Ca^2+^_MCH_ amplitudes across sleep states in the light phase (Fig. 2d,e). Moreover, drawing a threshold to divide the EEG Ratio trace into baseline versus active periods, we found that the correlation persisted during the active periods (Fig. 2d,f), which approximately matched onto REMS and transitional REMS periods (Fig. 2a,d). These results suggest that the correlation was not predominantly driven by baseline/non-active periods. Finally, the EEG Ratio led Ca^2+^_MCH_ by ∼ -1 to 1 sec (Fig. 2g) across individual recordings. Thus, empirically defined and optimized EEG Ratio tracks Ca^2+^_MCH_ in real time.

### EEG features extracted by machine-learning yielded similar correlations to Ca^2+^_MCH_

To minimize potential bias intrinsic to the empirical assay, we employed a machine-learning (ML)-based approach to assess whether Ca^2+^_MCH_ can be predicted by EEG signals. We calculated Z-scored time series of Ca^2+^_MCH_ signals and EEG power, and applied multi-layer perceptron (MLP) networks to predict Ca^2+^_MCH_ magnitude based on EEG power for the same time window. The correlation *r*-values between Ca^2+^_MCH_ and the EEG-predicted waveforms were used to assess the performance of ML. Based on a 10-fold cross-validation using data from pilot rats, we determined that networks with 3 hidden layers and 20 units/layer yielded best average performances (Methods; Fig. 3a,b). We then compared ML versus the empirical EEG Ratio (Fig. 3c-j). Both methods yielded similarly high *r*-values when applied to the training data set (Fig. 3c-f) or to new testing data sets (Fig. 3g). The performances of ML versus EEG Ratio on new testing data sets were similar and showed correlations, albeit with divergence in a subset of recordings (Fig. 3h). Overall, both ML-predicted waveforms and EEG Ratio were closely correlated with each other and with Ca^2+^_MCH_ (Fig. 3i-j). Examination of the trained ML model revealed that Layer 1 units placed consistent weights across *θ* and *δ* bands, often with opposite signs for the two frequency bands, whereas Layer 2 mostly responded to the difference of the two frequency bands (Fig. S2). Thus, MLP-based algorism and empirical EEG Ratio calculation similarly identified the *θ* - *δ* relationship as the key to predicting Ca^2+^_MCH_. We then chose the empirical EEG Ratio formula for all the following analyses, as it bears direct biological interpretations and requires less computational power for large sets of data processing.

### Consistent EEG-Ca^2+^_MCH_ correlations across sex, treatments, and light/dark phases

The EEG Ratio formula derived from the pilot male rats was then applied to additional male and female rats (Fig. 4a), which yielded similarly high *r*-values from recordings during the light phase (Fig. 4b), indicating high real-time correlations between EEG Ratio and Ca^2+^_MCH_ levels in rats of both sexes.

Does the correlation persist following cocaine experience, given that repeated cocaine use followed by withdrawal impairs MCH neuronal functions (21)? To address this, we measured EEG-Ca^2+^_MCH_ correlation in rats after long-term withdrawal from cocaine self-administration. Moreover, warming ambient temperature to close to thermoneutrality increases REMS in both drug-naïve and cocaine-experienced animals (22, 41), raising the question whether the correlation persists under warming-induced REMS changes. As shown in Fig. 4c, following either cocaine withdrawal and/or REMS interventions by environmental warming, the EEG-Ca^2+^_MCH_ correlation *r*-values remained high, and the slope of the linear regression was largely consistent (Fig. 4d). Furthermore, in response to REMS interventions such as sleep disturbance-rebound or warming-enhanced REMS, the EEG Ratio and Ca^2+^_MCH_ co-varied in individual rats linearly (Fig. 4e). These results indicate consistent correlations between EEG Ratio and Ca^2+^_MCH_ under different sleep conditions. Thus, during the light phase, when Ca^2+^_MCH_ predominantly occurs in REMS, EEG Ratio - Ca^2+^_MCH_ correlations are robust across sex and over diverse treatment conditions.

In the dark phase, the infrequent occurrence of REMS limited sampling of MCH neuron activities. However, MCH neurons are also activated in wakefulness during explorative behaviors with a delayed onset (23). Indeed, during rats’ explorative behaviors, Ca^2+^_MCH_ showed low-amplitude, asynchronous activities, which were accompanied by low-amplitude, asynchronous EEG Ratio events (shortened as “EEG events”, Fig. 4f). The linear regression *r*-values were lower under this condition compared to light phase (Fig. 4g-h). Occasionally, there were EEG events without matched Ca^2+^_MCH_ activities (Fig. 4f *arrows*). This may be in part due to the asynchronous nature of the Ca^2+^_MCH_ activities combined with the limited optic fiber size and light penetration for the fiber photometry recordings (22, 42) – i.e. there may be undetected Ca^2+^_MCH_ activities corresponding to the EEG events. Despite these few unmatched events, when randomly sampled across all ZT times of the dark phase in 2-h windows, the overall *r*-values remained high (Fig. 4i). Thus, it was feasible to use 24-h EEG Ratio measures to approximate MCH neuron activities across both light and dark phases.

### Cocaine-induced long-term changes in EEG Ratio

We previously reported that repeated cocaine self-administration and long-term withdrawal leads to hypo-functioning of MCH neurons via reduced membrane excitability and deficient glutamatergic transmissions (22). How, then, may cocaine-induced changes in MCH neurons be reflected in EEG Ratio measures? Rats were trained to self-administer cocaine (0.75 mg/kg/infusion at FR1, overnight + 2 h/d x 5 d; Fig. S3), and their sleep EEG was measured from before cocaine exposure and 21 days (d) after cocaine withdrawal (Fig. 5a). Comparing 24-h EEG Ratio features before cocaine exposure versus after 21-d withdrawal (Fig. 5b), there was a shift in the distributions of EEG events (defined as > “threshold”; threshold = 1.5 x amplitude of 60% dwell-time to eliminate baseline periods) – a decrease in total duration of EEG events in the dark phase and an increase in the area-under-curve in the light phase (Fig. 5c,d). How are these changes related to REMS measures? We reasoned that the EEG events’ total duration would approximately match the REMS duration, and their area-under-curve may be indicative of REMS intensity. During REMS, delta power is low (Fig. 1c), and REMS intensity is typically reflected in theta power. Thus, the conventional measure of REMS theta power x duration of REMS, which we defined as theta energy during REMS, may correspond to the area-under-curve of EEG Ratio. As shown in Fig. 5e,f, the shifts in EEG events were accompanied by decreases in REMS time and REMS *θ* energy in the dark phase and an increase in REMS *θ* energy in the light phase. Similarly, cluster analysis showed a decrease in average durations of EEG event-clusters in the dark phase (Fig. 5g). However, durations of long (>50 sec) EEG event-clusters were decreased in both dark and light phases on withdrawal d21 (Fig. 5h), suggesting a greater susceptibility of prolonged MCH neuron population activities to cocaine withdrawal. By contrast, REMS bout analysis did not detect changes in the average duration in dark or light phases (Fig. 5i), nor in long-REMS episodes in light phase (Fig. 5j). The above EEG Ratio changes were not likely because of the rats developing from young adulthood to adulthood, as age-matched control rats did not show changes in these EEG Ratio measures during the same ∼ 5 weeks of recordings (Fig. S4). These results suggest that EEG Ratio features are persistently altered following cocaine experience, reflecting REMS disturbance in the dark phase and deficient MCH neuron population activities in both dark and light phases. The results also demonstrate that EEG Ratio events represent a novel functional measure of REMS complementary to conventional REMS parameters with certain advantages.

### EEG Ratio features show predictive correlations with subsequent cocaine seeking behaviors

May certain EEG Ratio features correlate with cocaine intake? We first tested whether EEG Ratio features before cocaine exposure correlate with cocaine intake during the subsequent self-administration training (same rats as in Fig. 5a). In naïve rats, the relative distribution of EEG events in dark versus light phase before cocaine exposure was positively correlated with cocaine intake during subsequent self-administration training (Fig. 6a,b). Moreover, even when cocaine experience altered the overall dark-light distribution of EEG events (Fig. 5b,c), the correlations were preserved after withdrawal (Fig. 6c,d). By comparison, the corresponding REMS features from the same rats showed qualitatively similar, though less robust, correlations at both time points (Fig. 6e-h). Additionally, cluster analysis revealed that the EEG events’ average durations in light phase were inversely correlated with subsequent cocaine intake (Fig. 6i), and the inter-cluster intervals exhibited positive correlation (Fig. 6j). By comparison, the corresponding REMS parameters, average REMS bout durations, and intervals showed no trend or similar trend (Fig. 6k,l). Finally, long (>50 sec) clusters of EEG events from both before cocaine exposure and on withdrawal d21 consistently showed inverse correlations with cocaine intake during self-administration training (Fig. 6m,n). By comparison, long REMS average durations also showed similar correlations (Fig. 6o,p). These results suggest that the basal EEG Ratio and REMS features may predict future drug taking or seeking propensity.

**Figure 6.**
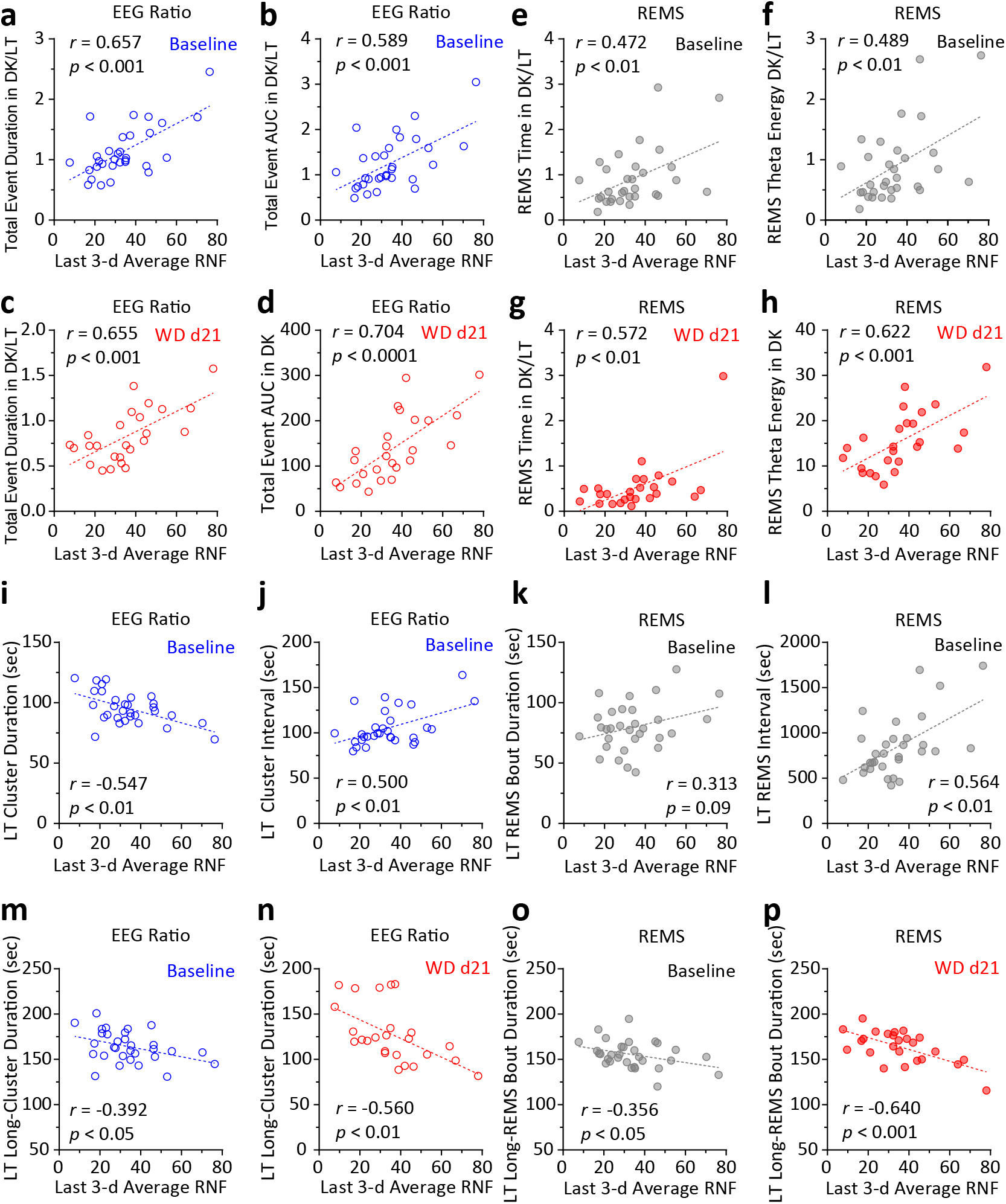
Baseline EEG Ratio features predict future cocaine intake. **a,b** Dark-light distribution ratio of EEG Ratio events at baseline before cocaine exposure showed correlations with cocaine intake during subsequent self-administration training. **c,d** Same EEG Ratio features at withdrawal d21 showed correlations with cocaine intake during self-administration training. **e-h** Corresponding REMS features from before cocaine exposure (**e,f**) or withdrawal d21 (**g,h**) showed correlations with cocaine intake during self-administration training. **i,j** Cluster analysis of EEG events showed that the average durations (**i**) or inter-cluster intervals (**j**) from before cocaine exposure were correlated with cocaine intake during self-administration training. **k,l** Corresponding REMS features from before cocaine exposure showed varying extent of correlations with cocaine intake during self-administration training. **m,n** Long (>50 sec)-EEG event clusters showed inverse correlations between average cluster durations and the amount of cocaine intake during self-administration training, both at baseline (**m**) and on withdrawal d21(**n**). **o,p** Long-REMS episodes showed inverse correlations between average bout durations and the amount of cocaine intake during self-administration training, both at baseline (**o**) and on withdrawal d21(**p**). RNF = reinforcement (i.e. cocaine infusions). Each circle represents a rat. n=31 male rats.

### EEG Ratio features correlate with incubation of cocaine craving

Relapse to drug use is a major challenge in treating SUD (43, 44). A prominent relapse rat model is the time-dependent intensification of cue-induced drug seeking after withdrawal from cocaine self-administration – termed “incubation” of drug craving – which indicates increased propensity for drug relapse after withdrawal (45–47). In a separate cohort of rats, we tested whether EEG Ratio features after long-term cocaine withdrawal are correlated with cocaine incubation. Rats underwent cocaine self-administration training similarly as before and long-term withdrawal up till d45, and sleep was measured ∼ WD d40-44 (Fig. 7a, S5). An incubation index was defined by the ratio of drug seeking (active nose-pokes, ANP) on withdrawal d45/d1, with an index >1 indicating incubation. Alternatively, change-of-ANP incubation was defined by ANP on withdrawal d45 - d1. The total area-under-curve of EEG events, which corresponds to the overall activity of MCH neurons over 24 h, exhibited a trending negative correlation with the incubation index (Fig. 7b) as well as the change-of-ANP incubation (Fig. 7c). This was consistent with the anti-relapse effects of MCH neuron activities that we recently reported (22). Moreover, cluster analysis of EEG events revealed that the 24-h total number of long clusters were inversely correlated with cocaine seeking on both withdrawal d1 and d45 (Fig. 7d,e), but not with incubation (data not shown). None of these EEG Ratio measures showed correlations with the mean cocaine intake during training. These results suggest that EEG Ratio parameters are differentially associated with different aspects of cocaine seeking behaviors. By comparison, the corresponding REMS measures from the same rats, such as total REMS *θ* energy and # of long REMS bouts, did not show similar correlations (Fig. 7f-i). Together, these results suggest that the EEG Ratio represents distinct features of REMS-related functions, and it provides unique quantifiable means to relate to behavioral outcomes.

**Figure 7.**
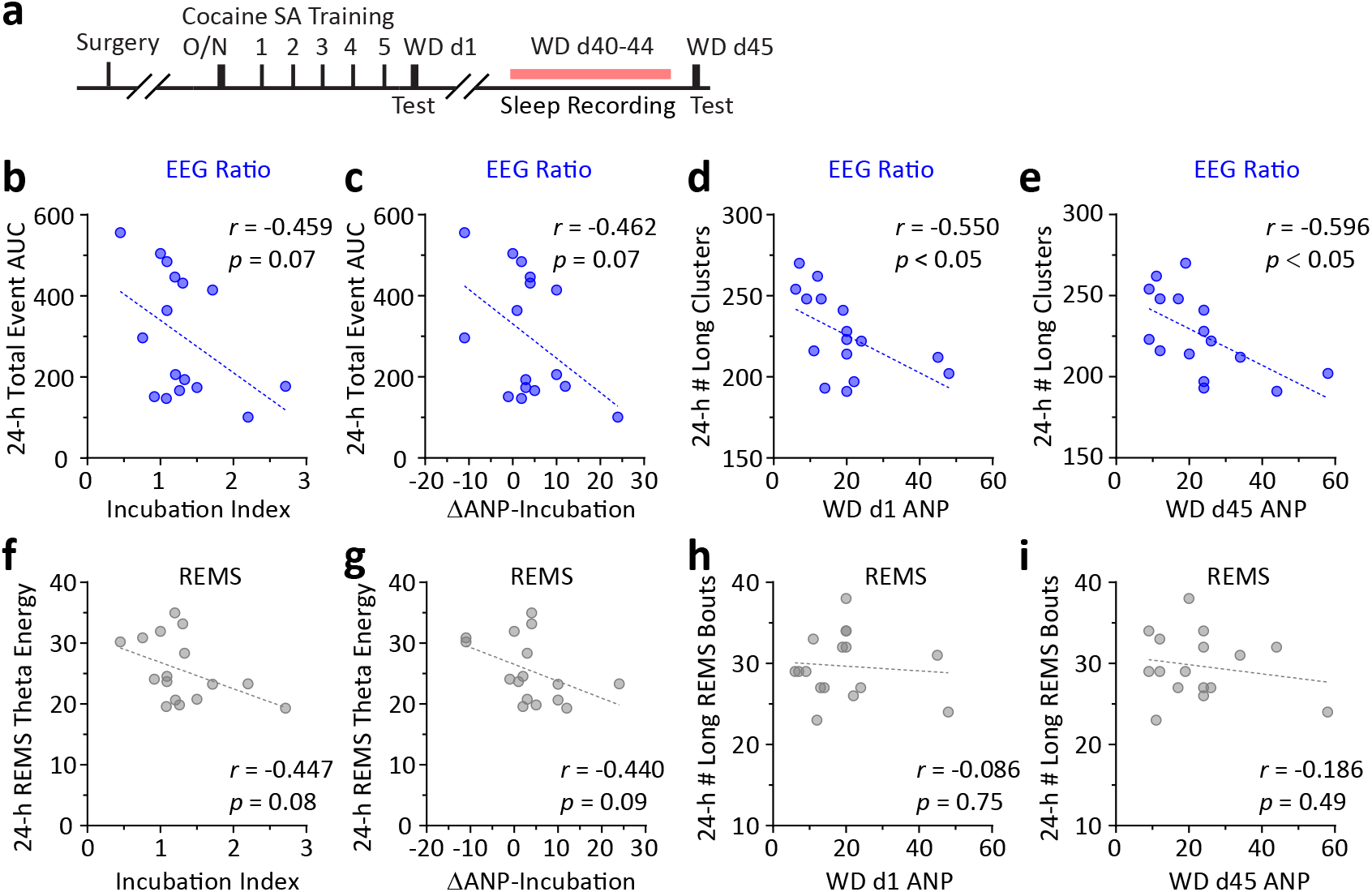
EEG Ratio features after long-term withdrawal are correlated with incubation of cocaine craving. **a** Cocaine self-administration training and EEG recording timeline. **b**,**c** 24-h EEG Ratio total event AUC was negatively correlated with the incubation index (**b**) and change-of-ANP (active nose-pokes on withdrawal d45-d1) incubation (**c**) on withdrawal d45. **d**,**e** Total # of EEG Ratio long clusters was negatively correlated with #ANP on withdrawal d1 (**d**) and on withdrawal d45 (**e**). **f**,**g** Corresponding REMS features such as 24-h total REMS *θ* energy did not show correlations with the incubation index (**f**) or change-of-ANP incubation (**g**) on withdrawal d45. **h**,**i** Total # of long REMS bouts did not show correlations with #ANP on withdrawal d1 (**h**) or on withdrawal d45 (**i**). Each circle represents a rat. n=16 male rats.

### Use EEG Ratio and REMS features to predict cocaine seeking

To what extent may EEG Ratio or REMS features predict future cocaine seeking? We focused on baseline sleep and EEG prior to cocaine exposure, using features highlighted in Figs. 6a,b,e,f. First, it was assumed that the outcome variable – amount of cocaine intake during self-administration training – was continuous, and the cross validated *r* were compared. The two EEG Ratio features showed better performances than the two REMS features (Table 1). Alternatively, the outcome (i.e. cocaine intake) was treated as an ordinal variable using 25% and 75% dividing lines, and Somers’ D and root-mean-square error (rMSE) were compared among the four features. Somers’D measures the categorical ranking consistency and rMSE evaluates the prediction error (48, 49). The REMS features showed better performances than the EEG Ratio features in categorically predicting levels of cocaine intake (Table 2). Together, these results suggest that EEG Ratio and REMS features show complementary strengths in predicting future cocaine seeking propensities.

**Table 1.** Evaluation of performance for predicting cocaine intake during SA training (continuous outcome)

|  | EEG Ratio |  | REMS |  |
| --- | --- | --- | --- | --- |
|  | Total Time | Total AUC | Total Time | Total Theta Energy |
| Predicted $r$ | 0.576 | 0.483 | 0.266 | 0.316 |
| Cross-validated $r$ | 0.657 | 0.590 | 0.472 | 0.489 |

**Table 2.** Evaluation of performance for predicting cocaine intake during SA training (ordinal outcome)

|  | EEG Ratio |  | REMS |  |
| --- | --- | --- | --- | --- |
|  | Total Time | Total AUC | Total Time | Total Theta Energy |
| Normalized rMSE std | 1.026 | 1.109 | 0.937 | 0.905 |
| Cross-validated Somers'D | 0.307 | 0.118 | 0.548 | 0.639 |

## Discussion

Activities in EEG frequency bands vary characteristically across sleep and wakefulness cycles. High *θ* activity is typically observed during REMS and certain wakefulness periods, and high *δ* activity during slow-wave sleep (50, 51). An increase in *θ*/*δ* ratio is associated with REMS and active wakefulness, thus can serve as one of the criteria for determining sleep states (39, 40). For this purpose, however, *θ*/*δ* ratio is typically calculated in 20-30 sec blocks, and rarely has it been analyzed at 1-sec resolution dynamics, nor has such dynamics been associated with specific cellular activities. Our characterization of a modified *θ*/*δ* ratio combines *θ* peak position shifts, and tracks LH MCH neuron activities at 1-sec resolution. This EEG Ratio represents a surface EEG signature for potential non-invasive monitoring of MCH neuron activities *in vivo*. This is also a first example of using EEG to deduce hypothalamic activities in a cell type-specific manner. The EEG Ratio described in this study also provides a unique functional measure of REMS, with metrics suitable for developing biomarkers.

Identifying biomarkers that predict future drug use propensity may open new avenues for individualized medicine in preventing and treating substance use disorders (SUDs) (1, 2). Although sleep dysregulation is often observed in chronic substance users that persists long after cessation of substance use (52), there is limited understanding of potential sleep biomarkers that may suggest future drug relapse propensity. Using rat cocaine self-administration model, our previous studies revealed an anti-relapse role of REMS and attempted to extract REMS features to predict SUD propensity (22). Similarly, in chronic alcohol users, REMS pressure, measured by REMS latency and density, at 2-week withdrawal is indicative of the likelihood to relapse within 6 months (53). However, it is not known the neural substrates underlying this, the accuracy of the predictions, or the applicability to other SUDs. Here we show that EEG Ratio clusters, reflecting LH MCH neuron activities and most prominent during REMS, have features correlate with future cocaine intake or seeking behaviors. It is important to note that our previous and current studies all used male rats to characterize REMS-drug use relationships (22, 27), and it is not known if similar behavioral predictions apply to female rats.

EEG Ratio events are closely related to REMS, including total time, cluster durations (versus REMS bout durations), and area-under-curve (versus REMS *θ* energy). The two also have important differences: i) EEG Ratio provides higher temporal resolution – at 1-sec – compared to REMS’s typical 10-sec (rodents) to 30-sec (human) resolution; ii) for REMS waveforms, EEG Ratio depicts the dynamics of *θ*/*δ* ratio together with *θ* peak position shifts within and across individual REMS bouts. These features are either omitted or grossly represented in the conventional measures of REMS intensity (i.e. averaged REMS *θ* power or energy); iii) EEG Ratio approximates LH MCH neuron population activities, whereas REMS waveforms predominantly reflect cortical and hippocampal activities. Therefore, EEG Ratio differs from, and complement, conventional REMS measures.

How may we use EEG Ratio to monitor MCH neuron activities *in vivo*? For example, a decrease in EEG events in the dark phase following cocaine experience (Fig. 5c,g,h) indicates decreased MCH neuron activities. On the other hand, inhibiting or eliminating MCH neurons does not typically alter REMS architecture (8, 54, 55). Thus, it remains to be determined whether an increase in EEG events necessarily indicates increased MCH neuron activities *in vivo*.

What are the biological interpretations of EEG Ratio features that predict future drug use? Taking advantage of the high throughput nature of EEG Ratio calculation independent of sleep scoring, we were able to analyze the 24-h EEG from a large number of rats before cocaine exposure, and identified EEG Ratio features that predict future drug intake propensity. Specifically, a larger dark-to-light ratio of total EEG event duration or area-under-curve predicted higher drug intake (Fig. 6a,b), and this correlation persisted following cocaine experience and withdrawal (Fig. 6c,d). Thus, cocaine-induced shifts in EEG event distributions from dark toward light phase may represent an adaptation to limit drug intake. Indeed, suppressing REMS in dark phase and allowing subsequent REMS rebound in light phase reduces craving-like behaviors in rats after withdrawal from cocaine self-administration (27). Additionally, the cluster-durations of the EEG events – which indicates consolidated REMS – both before (Fig. 6i,m) and after cocaine exposure (Fig. 6n) were negatively correlated with cocaine intake, and the number of long clusters of the EEG events was negatively correlated with cue-induced cocaine seeking after withdrawal (Fig. 7e). Related to these, long-bout REMS, which is the predominant REMS state that engages MCH neuron activities (22), was negatively correlated with cocaine intake (Fig. 6o,p). These results echo our previous findings that selectively increasing MCH neuron activities in sleep either by environmental warming or chemo/optogenetic stimulations of MCH neurons reduces craving-like behaviors in rats after cocaine withdrawal (22). Together, these results suggest that EEG Ratio or REMS features depicting prolonged synchronous activities of MCH neurons especially during the light phase may predict against future drug use.

May EEG Ratio characterized in the rat be translated across species? Given the universal nature of REMS being associated with the relative increase in *θ* and decrease in *δ* frequency powers, EEG Ratio calculation is likely feasible across species and reflects REMS. However, there are technical and species-specific features that need to be considered. For example, human EEGs are typically configured differently from the frontal-parietal differential EEG, there are significant slow wave activities during human REMS (56), the rodent-corresponding human REMS *θ* waves occur at lower frequencies (i.e., “slow *θ*”) (57–60), and REMS *θ* waves may also arise in different cortical and subcortical regions (57, 61–63). Nonetheless, human REMS exhibits significant *θ* activities in the prefrontal cortex (64), hippocampus (60, 63), and occipital cortex (65, 66), detectable in surface recordings (60); and human REMS *θ* is closely associated with emotion regulations (57, 67). Therefore, it is highly promising to translate EEG Ratio in humans.

The cellular and circuit mechanisms underlying coordinated changes that orchestrate EEG *θ* and *δ* fluctuations are not known. These changes may be independent of, but regulated by, MCH neuron activities, which in turn feed-back to REMS initiation and maintenance (8, 21, 54, 55). Given the importance of MCH neuron activities and MCH signaling in regulating mood and drug seeking (10, 22), these results provide a strong support to investigate EEG Ratio as a closely associated biometric in its potential for predicting future emotion and motivation-related behaviors. On the other hand, EEG Ratio features as biomarkers may exist independent of MCH neuron mechanisms. For example, EEG Ratio fluctuations may reflect or drive several parallel circuits to achieve overall REMS functions, and MCH neuron activity may be one of them. Thus, EEG Ratio features may predict behavioral outcomes through parallel circuit mechanisms not necessarily dependent on MCH neuron activities. In this sense, whether EEG Ratio calculated in different species accurately reflects MCH neuron activities in the particular species is not of significant concern when translating EEG Ratio across species.

## Acknowledgements

We thank Rachel L Hines, Braden R Bubarth, Bo Chen, and Nicholas J Lehman for assistance with rat behavioral training and testing; Myles Billard (Tucker-Davis Technologies) for fiber photometry technical support; Rachel L Hines, Braden R Bubarth, Peiran Zhou, Tyler R Barnhardt, Filip Hahs, and Baihe Zhang for assistance with data analyses.

## Funding

Research reported in this publication was supported by National Institutes of Health grant DA043826 (YH), DA046491 (YH), AA028145 (YH), DA057954 (YH), DA023206 (YD), DA040620 (YD), DA051010 (YD), DA047861 (YD), DA046346 (GT, YH), NS121913 (CH)

## Reagent

Cocaine was supplied by the Drug Supply Program of NIH NIDA.

## Author contributions

Conceptualization: YHH

Methodology: YHH, CH, GCT, RG, JW, SO, GA, JF

Investigation: YW, DL, JW, RG, RY, LC, CH, YHH

Supervision: YHH, CH, GCT

Writing—original draft: YHH, YW, CH, DL, GCT

Writing—review & editing: YHH, YD, CH, GCT, YW, RG, GA, JF

## Competing interests

Dr. Giancarlo Allocca is the creator of Somnivore software, and the owner of Somnivore Pty. Ltd. Dr. Jidong Fang is the creator of SleepMaster software, and the owner of Biosoft Studio. All other authors declare no conflict of interest. The content is solely the responsibility of the authors and does not necessarily represent the official views of the National Institutes of Health.

## Data and materials availability

All data needed to evaluate the conclusions in the paper are present in the paper and/or the Supplementary Materials.

## Supplementary Materials

**Figure S1.**
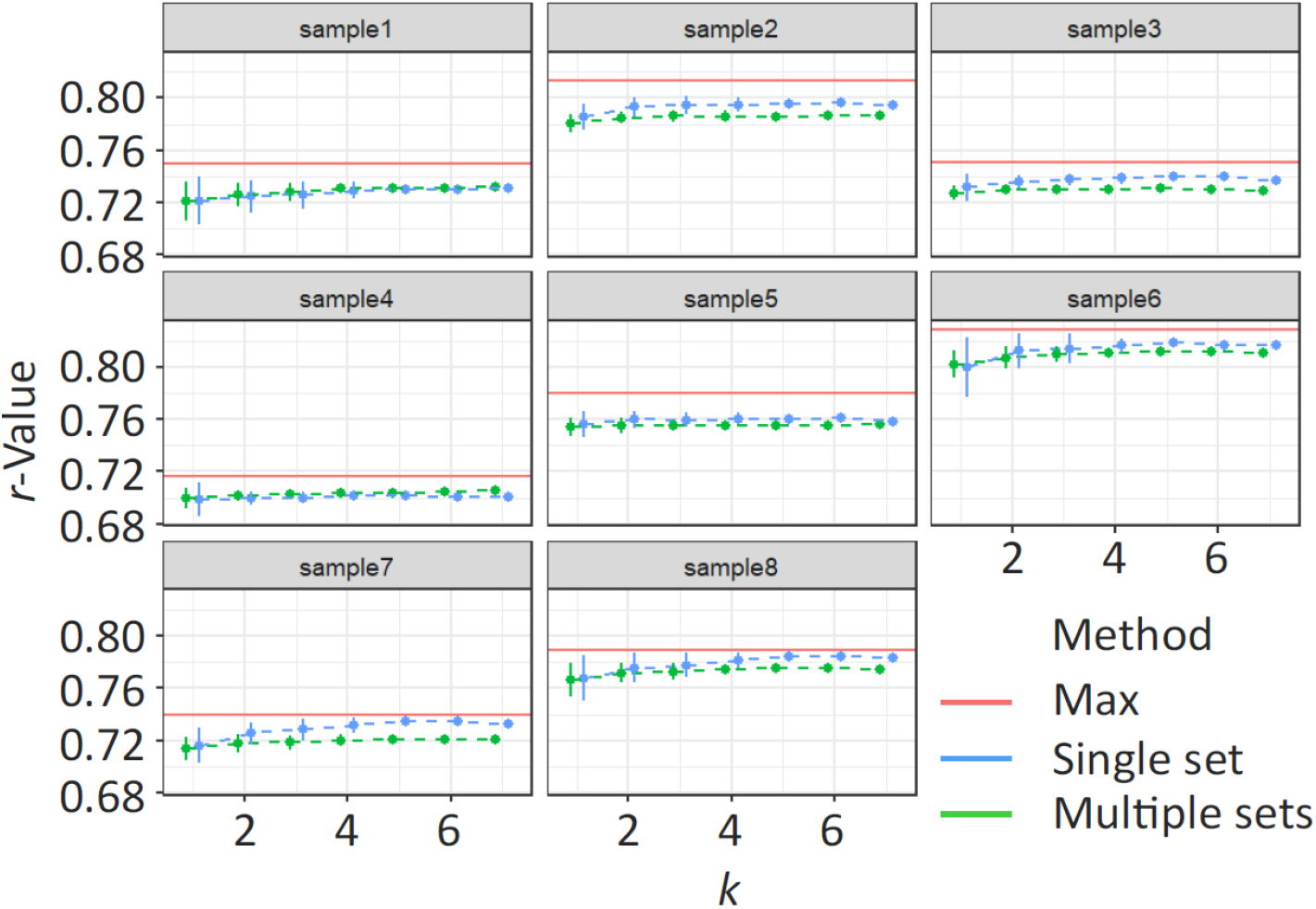
Comparing predictive performance using a single set of optimal parameters versus using multiple sets of best-performing parameters. *Red*: Maximal *r*-value achieved using a set of parameters optimized for the individual rat; *green*: *r*-values achieved using multiple sets of best-performing parameters across different rats; *blue*: *r*-values achieved using a single set of parameters (θ: 6.5-9.5 Hz, δ: 1-5.5 Hz) optimized across different rats. For cross-validation, *k* (*k* = 1,…,7) out of 8 rats were subsampled as the training set and the remaining rats as the test set. Mean +/- one standard deviation from the mean.

**Figure S2.**
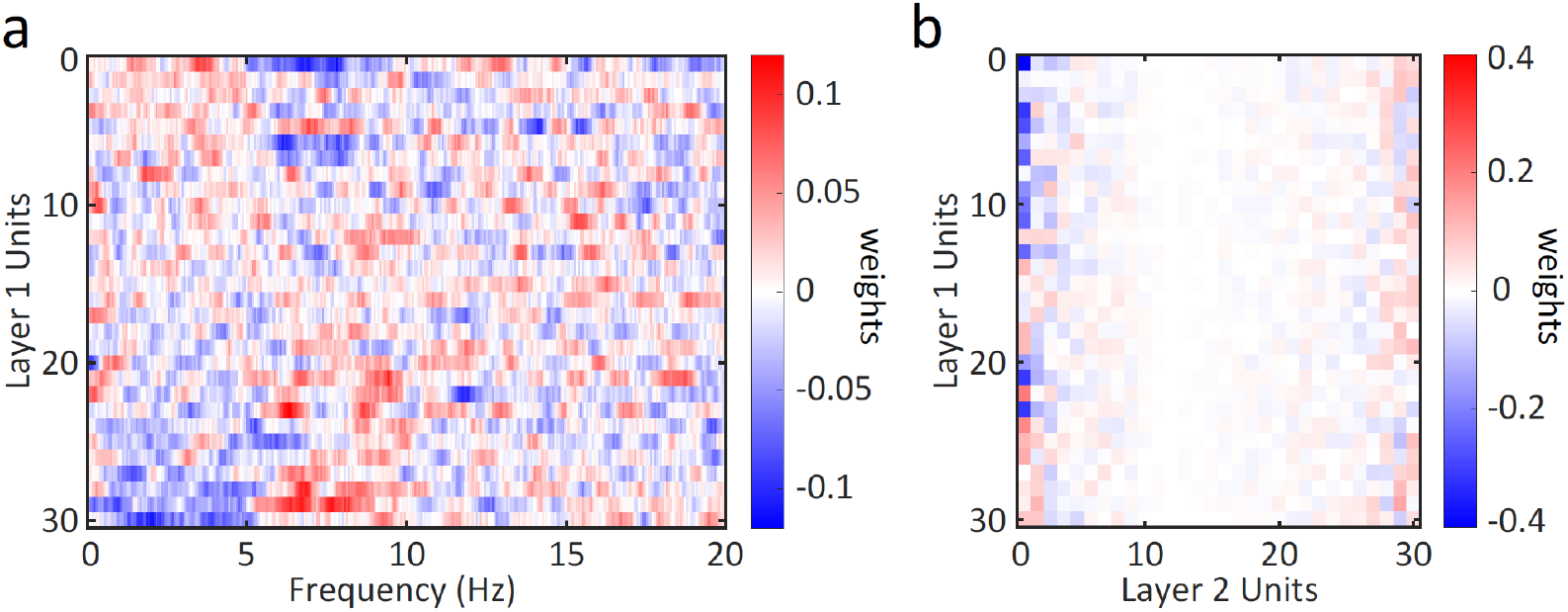
Weight structure of the trained ML model. **a** The weights to individual Layer 1 unit across frequency. Layer 1 units were sorted by the average weights between 1-5 Hz. The weights were smoothed using 0.5 Hz window. The units showed consistent weights across θ and δ bands, and weights tended to be opposite for these two frequency bands. **b** The weights from Layer 1 to Layer 2 units. Layer 2 units were sorted by their weights to the output unit with descending order. The first Layer 2 unit had the largest weight to the output unit. It had negative weights on the Layer 1 units that receive positive inputs from δ band, and mostly positive weights from Layer 1 units that receive more inputs from θ band. Hence it responded mostly to the power difference of the two frequency bands. Note that in addition to weights the bias terms and the nonlinear transfer functions in the ML model also affect model prediction.

**Figure S3.**
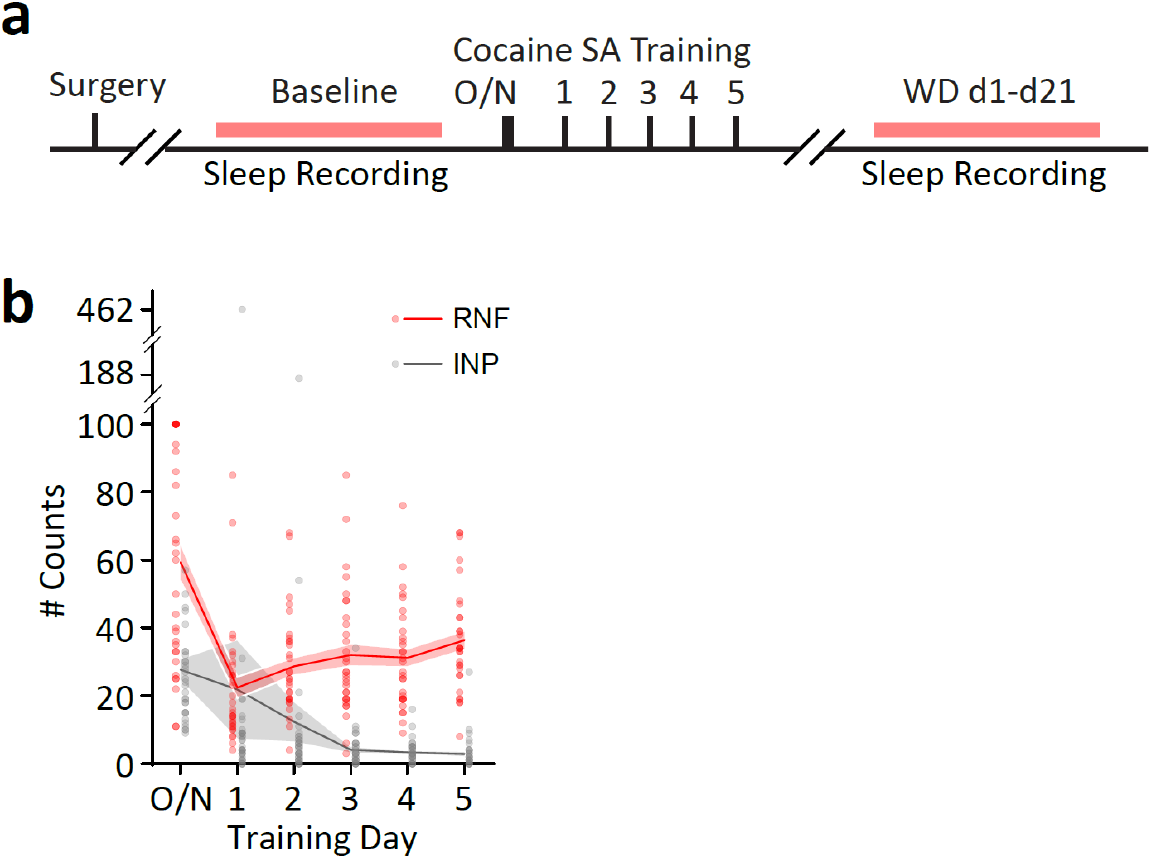
Cocaine self-administration training and withdrawal that complement Figures 5 and 6. **a** Timeline. **b** Cocaine self-administration training consisted of one overnight training followed by 5 days of 2-h daily training (0.75 mg/kg/infusion at fixed ratio 1). RNF = reinforcement (i.e. cocaine infusions). INP = inactive nose-pokes. Each circle represents a rat. n=31 male rats.

**Figure S4.**
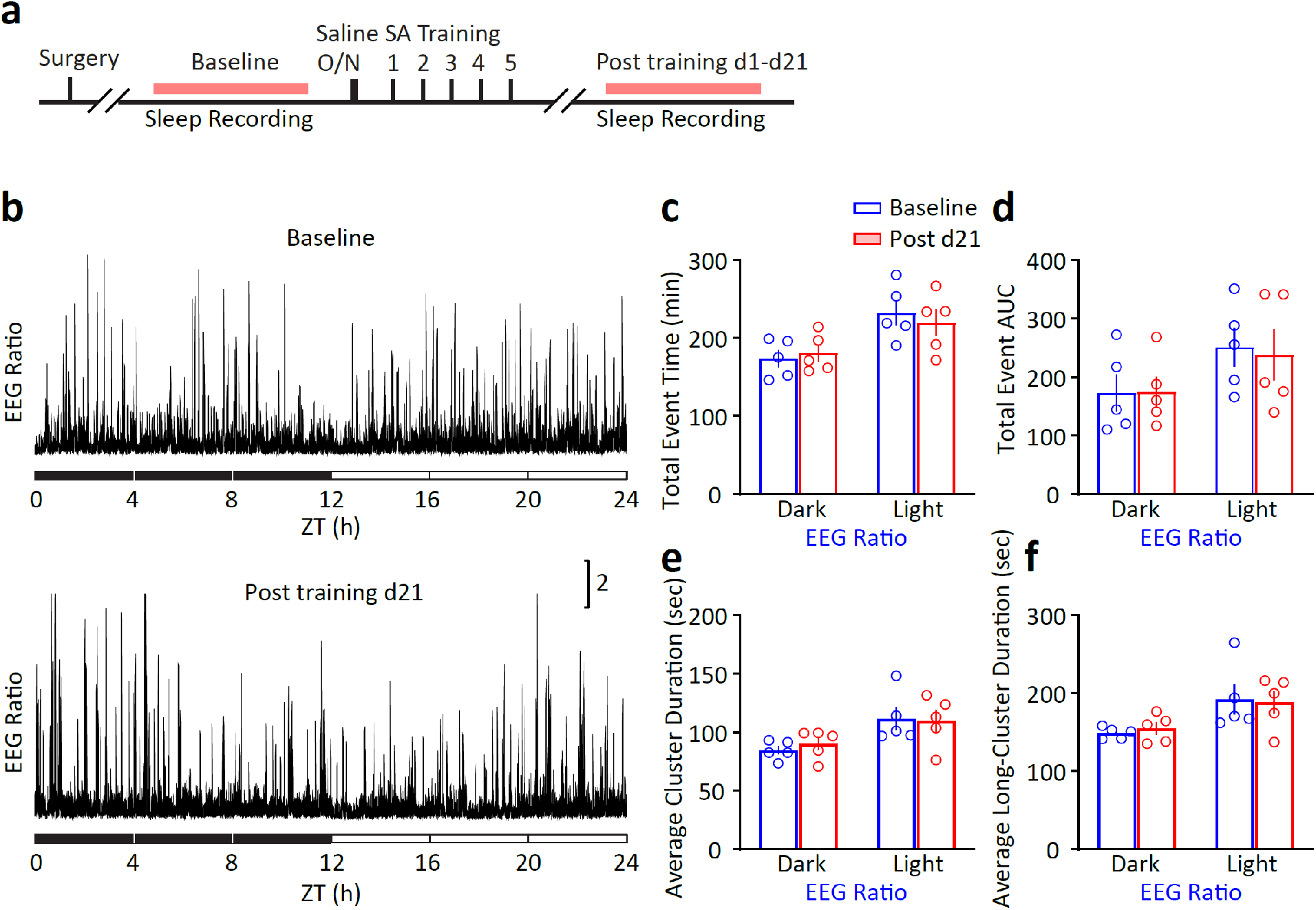
24-h EEG Ratio distribution does not change after withdrawal from saline self-administration. **a** Saline self-administration training and EEG recording timeline. **b** Example 24-h EEG Ratio extracted from baseline sleep before training (*top*) or 21 days after training (*bottom*). **c**,**d** The EEG Ratio distribution did not change over the dark or light phase in total event time (**c**) or total AUC (**d**). **c** interaction: F_1, 8_=0.251, *p*=0.630; **d** interaction: F_1, 8_=0.128, *p*=0.730; two-way RM ANOVA with Sidak *post-hoc* test. **e** Cluster analysis of EEG Ratio showed no change in the average cluster durations in the dark or light phase. **f** Long (>50 sec) EEG event clusters showed no change in durations in either dark or light phase. **e** interaction: F_1, 8_=0.176, *p*=0.686; **f** interaction: F_1, 8_=0.105, *p*=0.755; two-way RM ANOVA with Sidak *post-hoc* test.

**Figure S5.**
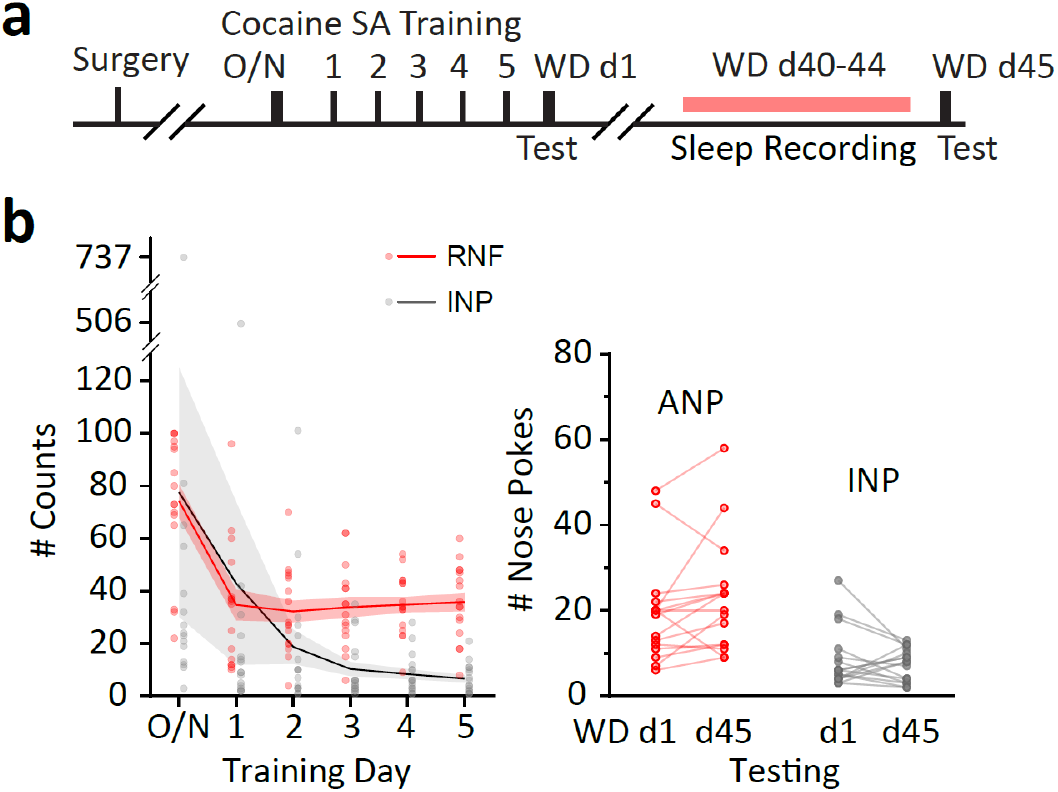
Cocaine self-administration training and withdrawal that complement Figure 7. **a** Timeline. **b** Cocaine self-administration training consisted of one overnight training followed by 5 days of 2-h daily training (0.75 mg/kg/infusion at fixed ratio 1). RNF = reinforcement (i.e. cocaine infusions). INP = inactive nose-pokes. Each circle represents a rat. n=16 male rats. Part of the data were used in (1).

## References

1. van der Stel J (2015): Precision in Addiction Care: Does It Make a Difference? Yale J Biol Med. 88:415–422.

2. Collins FS, Brown MK (2021): National Institutes of Health Sleep Research Plan. Advancing the Science of Sleep and Circadian Research.

3. Bonnavion P, Mickelsen LE, Fujita A, de Lecea L, Jackson AC (2016): Hubs and spokes of the lateral hypothalamus: cell types, circuits and behaviour. The Journal of physiology. 594:6443–6462.

4. Sharpe MJ (2023): The cognitive (lateral) hypothalamus. Trends Cogn Sci.

5. Elmquist JK, Elias CF, Saper CB (1999): From lesions to leptin: hypothalamic control of food intake and body weight. Neuron. 22:221–232.

6. Petrovich GD (2018): Lateral Hypothalamus as a Motivation-Cognition Interface in the Control of Feeding Behavior. Front Syst Neurosci. 12:14.

7. Bittencourt JC, Presse F, Arias C, Peto C, Vaughan J, Nahon JL, et al. (1992): The melanin-concentrating hormone system of the rat brain: an immuno- and hybridization histochemical characterization. The Journal of comparative neurology. 319:218–245.

8. Jego S, Glasgow SD, Herrera CG, Ekstrand M, Reed SJ, Boyce R, et al. (2013): Optogenetic identification of a rapid eye movement sleep modulatory circuit in the hypothalamus. Nat Neurosci. 16:1637–1643.

9. Lagos P, Torterolo P, Jantos H, Monti JM (2011): Immunoneutralization of melanin-concentrating hormone (MCH) in the dorsal raphe nucleus: effects on sleep and wakefulness. Brain research. 1369:112–118.

10. Torterolo P, Scorza C, Lagos P, Urbanavicius J, Benedetto L, Pascovich C, et al. (2015): Melanin-Concentrating Hormone (MCH): Role in REM Sleep and Depression. Front Neurosci. 9:475.

11. Al-Massadi O, Dieguez C, Schneeberger M, Lopez M, Schwaninger M, Prevot V, et al. (2021): Multifaceted actions of melanin-concentrating hormone on mammalian energy homeostasis. Nat Rev Endocrinol. 17:745–755.

12. Blasiak A, Gundlach AL, Hess G, Lewandowski MH (2017): Interactions of Circadian Rhythmicity, Stress and Orexigenic Neuropeptide Systems: Implications for Food Intake Control. Front Neurosci. 11:127.

13. Borowsky B, Durkin MM, Ogozalek K, Marzabadi MR, DeLeon J, Lagu B, et al. (2002): Antidepressant, anxiolytic and anorectic effects of a melanin-concentrating hormone-1 receptor antagonist. Nature medicine. 8:825–830.

14. Chung S, Parks GS, Lee C, Civelli O (2011): Recent updates on the melanin-concentrating hormone (MCH) and its receptor system: lessons from MCH1R antagonists. Journal of molecular neuroscience: MN. 43:115–121.

15. Macneil DJ (2013): The role of melanin-concentrating hormone and its receptors in energy homeostasis. Front Endocrinol (Lausanne). 4:49.

16. Haemmerle CA, Campos AM, Bittencourt JC (2015): Melanin-concentrating hormone inputs to the nucleus accumbens originate from distinct hypothalamic sources and are apposed to GABAergic and cholinergic cells in the Long-Evans rat brain. Neuroscience. 289:392–405.

17. Hassani OK, Lee MG, Jones BE (2009): Melanin-concentrating hormone neurons discharge in a reciprocal manner to orexin neurons across the sleep-wake cycle. Proceedings of the National Academy of Sciences of the United States of America. 106:2418–2422.

18. Izawa S, Chowdhury S, Miyazaki T, Mukai Y, Ono D, Inoue R, et al. (2019): REM sleep-active MCH neurons are involved in forgetting hippocampus-dependent memories. Science. 365:1308–1313.

19. Landmann N, Kuhn M, Maier JG, Spiegelhalder K, Baglioni C, Frase L, et al. (2015): REM sleep and memory reorganization: Potential relevance for psychiatry and psychotherapy. Neurobiol Learn Mem. 122:28–40.

20. Krause AJ, Simon EB, Mander BA, Greer SM, Saletin JM, Goldstein-Piekarski AN, et al. (2017): The sleep-deprived human brain. Nat Rev Neurosci. 18:404–418.

21. Wang Y, Guo R, Chen B, Rahman T, Cai L, Li Y, et al. (2020): Cocaine-induced neural adaptations in the lateral hypothalamic melanin-concentrating hormone neurons and the role in regulating rapid eye movement sleep after withdrawal. Mol Psychiatry.

22. Guo R, Wang Y, Yan R, Chen B, Ding W, Gorczyca MT, et al. (2022): REM Sleep Engages MCH Neurons to Reduce Cocaine Seeking. Biological psychiatry.

23. Blanco-Centurion C, Luo S, Spergel DJ, Vidal-Ortiz A, Oprisan SA, Van den Pol AN, et al. (2019): Dynamic Network Activation of Hypothalamic MCH Neurons in REM Sleep and Exploratory Behavior. The Journal of neuroscience: the official journal of the Society for Neuroscience. 39:4986–4998.

24. Teplan M (2002): Fundamentals of EEG Measurement. IEEE Measurement Science Review. 2:1–11.

25. Seeber M, Cantonas LM, Hoevels M, Sesia T, Visser-Vandewalle V, Michel CM (2019): Subcortical electrophysiological activity is detectable with high-density EEG source imaging. Nat Commun. 10:753.

26. Attal Y, Schwartz D (2013): Assessment of subcortical source localization using deep brain activity imaging model with minimum norm operators: a MEG study. PloS one. 8:e59856.

27. Chen B, Wang Y, Liu X, Liu Z, Dong Y, Huang YH (2015): Sleep Regulates Incubation of Cocaine Craving. The Journal of neuroscience: the official journal of the Society for Neuroscience. 35:13300–13310.

28. Ma YY, Lee BR, Wang X, Guo C, Liu L, Cui R, et al. (2014): Bidirectional modulation of incubation of cocaine craving by silent synapse-based remodeling of prefrontal cortex to accumbens projections. Neuron. 83:1453–1467.

29. Ma YY, Wang X, Huang Y, Marie H, Nestler EJ, Schluter OM, et al. (2016): Re-silencing of silent synapses unmasks anti-relapse effects of environmental enrichment. Proceedings of the National Academy of Sciences of the United States of America. 113:5089–5094.

30. Krueger JM, Obal F (1993): A neuronal group theory of sleep function. Journal of sleep research. 2:63–69.

31. Winters BD, Huang YH, Dong Y, Krueger JM (2011): Sleep loss alters synaptic and intrinsic neuronal properties in mouse prefrontal cortex. Brain research. 1420:1–7.

32. Liu Z, Wang Y, Cai L, Li Y, Chen B, Dong Y, et al. (2016): Prefrontal Cortex to Accumbens Projections in Sleep Regulation of Reward. The Journal of neuroscience: the official journal of the Society for Neuroscience. 36:7897–7910.

33. Lee BR, Ma YY, Huang YH, Wang X, Otaka M, Ishikawa M, et al. (2013): Maturation of silent synapses in amygdala-accumbens projection contributes to incubation of cocaine craving. Nat Neurosci. 16:1644–1651.

34. Ma YY, Lee BR, Wang X, Guo C, Liu L, Cui R, et al. (2014): Bidirectional Modulation of Incubation of Cocaine Craving by Silent Synapse-Based Remodeling of Prefrontal Cortex to Accumbens Projections. Neuron.

35. Allocca G, Ma S, Martelli D, Cerri M, Del Vecchio F, Bastianini S, et al. (2019): Validation of ‘Somnivore’, a Machine Learning Algorithm for Automated Scoring and Analysis of Polysomnography Data. Front Neurosci. 13:207.

36. Polyak BT (1964): Some methods of speeding up the convergence of iteration methods. USSR Computational Mathematics and Mathematical Physics. 4:17.

37. LeCun Y, Bengio Y, Hinton G (2015): Deep learning. Nature. 521:436–444.

38. Ghasemi A, Zahediasl S (2012): Normality tests for statistical analysis: a guide for non-statisticians. Int J Endocrinol Metab. 10:486–489.

39. Maloney KJ, Cape EG, Gotman J, Jones BE (1997): High-frequency gamma electroencephalogram activity in association with sleep-wake states and spontaneous behaviors in the rat. Neuroscience. 76:541–555.

40. Rayan A, Agarwal A, Samanta A, Severijnen E, van der Meij J, Genzel L (2024): Sleep scoring in rodents: Criteria, automatic approaches and outstanding issues. The European journal of neuroscience. 59:526–553.

41. Komagata N, Latifi B, Rusterholz T, Bassetti CLA, Adamantidis A, Schmidt MH (2019): Dynamic REM Sleep Modulation by Ambient Temperature and the Critical Role of the Melanin-Concentrating Hormone System. Current biology: CB. 29:1976–1987 e1974.

42. Yizhar O, Fenno LE, Davidson TJ, Mogri M, Deisseroth K (2011): Optogenetics in neural systems. Neuron. 71:9–34.

43. Hunt WA, Barnett LW, Branch LG (1971): Relapse rates in addiction programs. Journal of clinical psychology. 27:455–456.

44. McLellan AT, Lewis DC, O’Brien CP, Kleber HD (2000): Drug dependence, a chronic medical illness: implications for treatment, insurance, and outcomes evaluation. JAMA. 284:1689–1695.

45. Grimm JW, Hope BT, Wise RA, Shaham Y (2001): Neuroadaptation. Incubation of cocaine craving after withdrawal. Nature. 412:141–142.

46. Pickens CL, Airavaara M, Theberge F, Fanous S, Hope BT, Shaham Y (2011): Neurobiology of the incubation of drug craving. Trends in neurosciences. 34:411–420.

47. Tran-Nguyen LT, Fuchs RA, Coffey GP, Baker DA, O’Dell LE, Neisewander JL (1998): Time-dependent changes in cocaine-seeking behavior and extracellular dopamine levels in the amygdala during cocaine withdrawal. Neuropsychopharmacology: official publication of the American College of Neuropsychopharmacology. 19:48–59.

48. Newson R (2006): Confidence Intervals for Rank Statistics: Somers’ D and Extensions. The Stata Journal. 6:309–334.

49. Pham BT, Son LH, Hoang T-A, Nguyen D-M, Bui DT (2018): Prediction of shear strength of soft soil using machine learning methods. CATENA. 166:181–191.

50. Chellappa SL, Schmidt C, Cajochen C (2014): Neurophysiology of Sleep and Wakefulness. Sleepiness and Human Impact Assessment: Springer, pp 23–41.

51. Purves D, Augustine GJ, Fitzpatrick D, Katz LC, LaMantia A-S, McNamara JO, et al. (2001): Neuroscience, Chapter 28. Sleep and Wakefulness. Sunderland (MA): Sinauer Associates.

52. Angarita GA, Emadi N, Hodges S, Morgan PT (2016): Sleep abnormalities associated with alcohol, cannabis, cocaine, and opiate use: a comprehensive review. Addict Sci Clin Pract. 11:9.

53. Gann H, Feige B, Hohagen F, van Calker D, Geiss D, Dieter R (2001): Sleep and the cholinergic rapid eye movement sleep induction test in patients with primary alcohol dependence. Biological psychiatry. 50:383–390.

54. Konadhode RR, Pelluru D, Blanco-Centurion C, Zayachkivsky A, Liu M, Uhde T, et al. (2013): Optogenetic stimulation of MCH neurons increases sleep. The Journal of neuroscience: the official journal of the Society for Neuroscience. 33:10257–10263.

55. Tsunematsu T, Ueno T, Tabuchi S, Inutsuka A, Tanaka KF, Hasuwa H, et al. (2014): Optogenetic manipulation of activity and temporally controlled cell-specific ablation reveal a role for MCH neurons in sleep/wake regulation. The Journal of neuroscience: the official journal of the Society for Neuroscience. 34:6896–6909.

56. Bray N (2019): REM sleep makes slow waves. Nat Rev Neurosci. 20:191.

57. Hutchison IC, Rathore S (2015): The role of REM sleep theta activity in emotional memory. Front Psychol. 6:1439.

58. Moroni F, Nobili L, Curcio G, De Carli F, Fratello F, Marzano C, et al. (2007): Sleep in the human hippocampus: a stereo-EEG study. PloS one. 2:e867.

59. Lega BC, Jacobs J, Kahana M (2012): Human hippocampal theta oscillations and the formation of episodic memories. Hippocampus. 22:748–761.

60. Bodizs R, Kantor S, Szabo G, Szucs A, Eross L, Halasz P (2001): Rhythmic hippocampal slow oscillation characterizes REM sleep in humans. Hippocampus. 11:747–753.

61. Dong Y, Li J, Zhou M, Du Y, Liu D (2022): Cortical regulation of two-stage rapid eye movement sleep. Nat Neurosci. 25:1675–1682.

62. Wang Z, Fei X, Liu X, Wang Y, Hu Y, Peng W, et al. (2022): REM sleep is associated with distinct global cortical dynamics and controlled by occipital cortex. Nat Commun. 13:6896.

63. Cantero JL, Atienza M, Stickgold R, Kahana MJ, Madsen JR, Kocsis B (2003): Sleep-dependent theta oscillations in the human hippocampus and neocortex. J Neurosci. 23:10897–10903.

64. Vijayan S, Lepage KQ, Kopell NJ, Cash SS (2017): Frontal beta-theta network during REM sleep. eLife. 6.

65. Olejarczyk E, Gotman J, Frauscher B (2022): Region-specific complexity of the intracranial EEG in the sleeping human brain. Sci Rep. 12:451.

66. Frauscher B, Joshi S, von Ellenrieder N, Nguyen DK, Dubeau F, Gotman J (2018): Sharply contoured theta waves are the human correlate of ponto-geniculo-occipital waves in the primary visual cortex. Clin Neurophysiol. 129:1526–1533.

67. Cowdin N, Kobayashi I, Mellman TA (2014): Theta frequency activity during rapid eye movement (REM) sleep is greater in people with resilience versus PTSD. Exp Brain Res. 232:1479–1485.

## References

1. Guo R, Wang Y, Yan R, Chen B, Ding W, Gorczyca MT, et al. (2022): REM Sleep Engages MCH Neurons to Reduce Cocaine Seeking. Biological psychiatry.

